# Phagocytic “teeth” and myosin-II “jaw” power target constriction during phagocytosis

**DOI:** 10.1101/2021.03.14.435346

**Authors:** Daan Vorselen, Sarah R. Barger, Yifan Wang, Wei Cai, Julie A. Theriot, Nils C. Gauthier, Mira Krendel

**Author notes:** These two authors contributed equally (Daan Vorselen, Sarah R. Barger).

## Abstract

Phagocytosis requires rapid actin reorganization and spatially controlled force generation to ingest targets ranging from pathogens to apoptotic cells. How actomyosin activity directs membrane extensions to engulf such diverse targets remains unclear. Here, we combine lattice light-sheet microscopy (LLSM) with microparticle traction force microscopy (MP-TFM) to quantify actin dynamics and subcellular forces during macrophage phagocytosis. We show that spatially localized forces leading to target constriction are prominent during phagocytosis of antibody-opsonized targets. This constriction is largely mediated by Arp2/3-mediated assembly of discrete actin protrusions containing myosin 1e and 1f (“teeth”) that are interconnected in a ring-like organization. Contractile myosin-II activity contributes to late-stage phagocytic force generation and progression, suggesting a specific role in phagocytic cup closure. Observations of partial target eating attempts and sudden target release via a popping mechanism suggest that constriction may be critical for resolving complex *in vivo* target encounters. Overall, our findings suggest a phagocytic cup-shaping mechanism that is distinct from cytoskeletal remodeling in 2D cell motility and may contribute to mechanosensing and phagocytic plasticity.

## Introduction

Phagocytic uptake of microbial pathogens, apoptotic cells, and debris are essential processes for human health^1,2^. Given the variety of phagocytic targets, ranging widely in shape, size and mechanical stiffness, this process requires remarkable plasticity. Phagocytosis is initiated when phagocytic receptors recognize distinct molecular patterns coating the target^3,4^. Downstream signalling from phagocytic receptors then leads to the formation of membrane protrusions, guided around the target through sequential ligand engagement in a zipper-like fashion^5–7^, to make the phagocytic cup. Mechanically, progression of the phagocytic cup is powered by F-actin polymerization that pushes the plasma membrane forward and culminates in the eventual closure of the cup and the formation of a membrane-enclosed phagosome^8^.

Previously, F-actin within the phagocytic cup was generally considered to be homogenous and the membrane extensions around the target were frequently likened to lamellipodia at the leading edge of a migrating cell^7,9–11^. Yet recent studies have identified dynamic adhesions or podosome-like structures within the phagocytic cup that appear to be sites of actin polymerization^12,13^.

Podosomes are specialized F-actin adhesive structures that are prominent in myeloid cells and capable of generating traction forces and degrading extracellular matrix^14,15^. This suggests a fundamentally different mechanism for cup shaping (assembly of individual actin-based protrusions vs. a uniform actin meshwork), and since podosomes are mechanosensitive^15–17^, they may contribute to the mechanosensitivity observed in phagocytosis, whereby phagocytes engulf stiffer targets more readily than softer ones^8,18–21^.

Moreover, the mechanism of cup closure and the involvement of myosin motor proteins therein has remained elusive. Based on observations of phagocytes deforming red blood cells during internalization^22–24^, it has been hypothesized that closure of the phagocytic cup involves myosin-II mediated “purse-string” contractility^25,26^ However, the limited effect of myosin-II inhibition on phagocytic uptake efficiency^27,28^ and on traction forces tangential to the target surface has led to the view that myosin-II likely does not contribute to phagocytic internalization^7^.

It is clear that phagocytosis is driven by mechanical forces, but examining these forces has been challenging due to the limitations of experimental approaches. Changes in cellular tension during phagocytosis measured by micropipette aspiration have been well studied^29,30^, but this technique fails to capture cell-target forces and the spatial variation within the phagocytic cup. Recently, traction force microscopy (TFM) combined with a 2D spreading assay known as frustrated phagocytosis has been used to measure cell-target forces^12,20,25,31^, however, this assay fails to capture the biologically relevant geometry of the phagocytic cup, which very likely affects cytoskeletal dynamics and force generation. Moreover, TFM only measures forces tangential to the target surface, neglecting forces normal to the target surface, which may be critical. With the recent introduction of particle-based force sensing methods^32,33^, particularly microparticle traction force microscopy (MP-TFM)^21^, both normal and shear force components can now be studied throughout phagocytosis.

Here, we utilize live-cell imaging combining MP-TFM and lattice light-sheet microscopy (LLSM) to reveal how mechanical forces generated by actin polymerization and myosin contractility drive phagocytic engulfment mediated by Fc receptors (FcRs). We show that phagocytes assemble F-actin “teeth” that mediate target constriction throughout phagocytosis. Analysis of forces shows a unique signature, in which target constriction, or squeezing, is balanced by pulling forces at the base of the phagocytic cup at early stages and target compression throughout the cup at later stages. Together, normal forces far exceed shear forces at the cell-target interface, pointing to a mechanism fundamentally distinct from lamellipodial spreading in cell motility. We find that target constriction is mediated by Arp2/3-mediated actin polymerization throughout phagocytosis.

Moreover, based on both force analysis and precise quantitative measurement of cup progression, we establish a clear role for myosin-II purse string contractility, specifically in phagocytic cup closure. Finally, we present how this force signature might be critical for target selection and ingestion in more complex physiological settings.

## Results

### Lattice light-sheet microscopy (LLSM) reveals sequence of target deformations induced during live phagocytic engulfment

Given the fast, 3D and light-sensitive nature of phagocytosis, we used lattice light-sheet microscopy (LLSM) for high-speed volumetric imaging with minimal phototoxicity to investigate cytoskeletal dynamics and phagocytic forces. To monitor internalization in real time, RAW264.7 macrophages were transfected with mEmerald-Lifeact for labelling of filamentous actin and were fed deformable acrylamide-co-acrylic acid-microparticles (DAAMPs) (Fig. 1a). To investigate FcR-mediated phagocytosis, DAAMPs were functionalized with BSA and anti-BSA IgG, as well as AF647-Cadaverine for visualization^21^ (Fig. 1b). RAW macrophages fed DAAMPs with a diameter of 9 μm and a Young’s modulus of 1.4 or 6.5 kPa typically formed a chalice-shaped phagocytic cup and completed phagocytosis in similar timeframe (~3 minutes) as previously reported for polystyrene particles^34^ (Fig. 1a, Supplementary Video 1, Supplementary Fig. 1e). 3D shape reconstructions of the DAAMPs enabled us to examine target deformations as a direct readout of phagocytic forces in real time (Fig. 1b-c). Interestingly, we observed target constriction defined by discrete spots of deformation that appeared around the circumference of the DAAMP at the rim of the phagocytic cup (Fig. 1d, Supplementary Video 2). While these deformations were more apparent using the softer 1.4 kPa DAAMPs, the same force pattern was observed using stiffer 6.5 kPa targets (Supplementary Fig. 1a-b, Supplementary Video 3). These indentations travelled parallel to the direction of phagocytic cup elongation, along the length of the DAAM particle until cup closure and were associated with ~400 nm maximum target constriction for 1.4 kPa DAAMPs (Supplementary Fig. 1c). In addition, we observed bulk compression during phagocytic cup progression (~ 0.5 kPa) and after complete internalization of the DAAMPs (up to 1.5 kPa), leading to a dramatic reduction in DAAMP diameter (Supplementary Fig. 1f-h). The spherical appearance of targets and the gradual monotonic increase in compression after completion of engulfment suggests that this compression may relate to the recently observed shrinkage of internalized macropinosomes by osmotic pressure changes regulated by ion flux^35^. However, we sometimes observed the appearance of an F-actin shell around the target, similar to previous reports^36^, suggesting that cytoskeletal forces may also contribute to such compression (Supplementary Fig. 1i, Supplementary Video 4).

**Figure 1.**
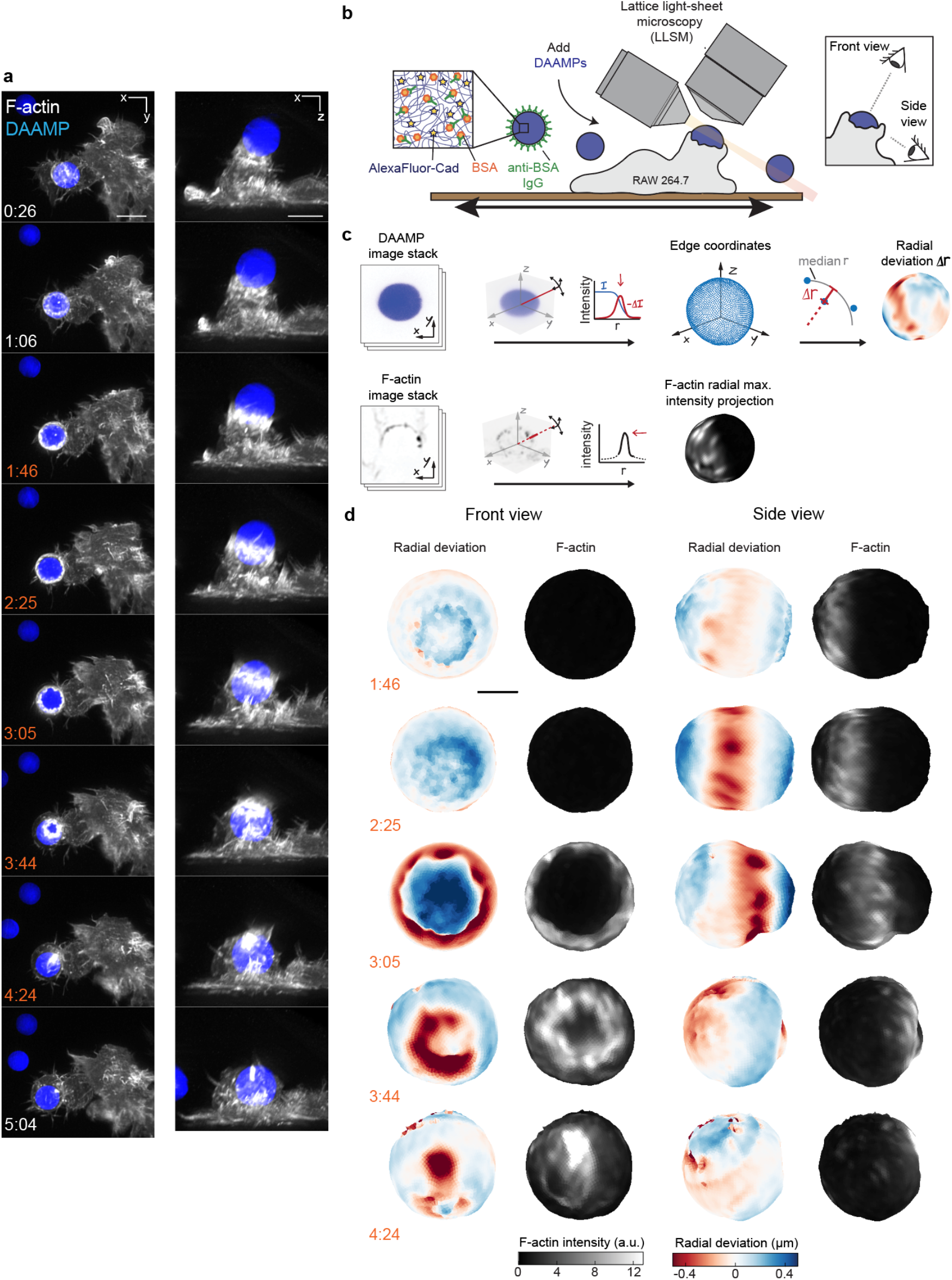
Combined MP-TFM and LLSM reveals phagocytic force-induced deformations in real time. RAW macrophages transfected with mEmerald-Lifeact were fed DAAM-particles (9μm,1.4 kPa) functionalized with AF647-Cadaverine, BSA and anti-BSA IgG and imaged using lattice light-sheet microscopy (LLSM). **a,** Time lapse montage (min:s) of maximum intensity projections in x/y and x/z. Scale bar, 5 μm **b,c** Schematic of the combined LLSM and MP-TFM experimental approach and analysis, respectively. **d,** Front and side view of reconstructed DAAM-particle internalized in **a** showing target deformations and F-actin localization on particle surface. Colorscale represents the deviation of each vertex from a perfect sphere with radius equal to the median radial distance of all edge coordinates to the particle centroid. Scale bar, 3 μm.

### Forces normal to the target surface are dominant during phagocytic engulfment and lead to strong target constriction

To more closely investigate the role of the actin cytoskeleton in force generation during FcR-mediated engulfment, we performed MP-TFM measurements on cells fixed during the process of phagocytosis. RAW macrophages were exposed to 9 μm, 1.4 kPa DAAMPs, after which they were fixed and stained for F-actin (Fig. 2a). Immunostaining of the exposed particle surface allowed precise determination of the stage of engulfment (Supplementary Fig. 2a). Confocal Z-stacks of phagocytic cups enabled 3D target shape reconstructions with super-resolution accuracy and inference of cellular traction forces (Fig. 2b,c). The deformation patterns and magnitudes for the fixed samples were similar to those observed in living cells by the LLSM imaging of live cells (Fig. 2b, Supplementary Fig. 1d & 2b). Specifically, we noted a ring of inhomogeneous F-actin localized along the rim of the phagocytic cup, where high F-actin intensity strongly correlated with inward target deformation.

**Figure 2.**
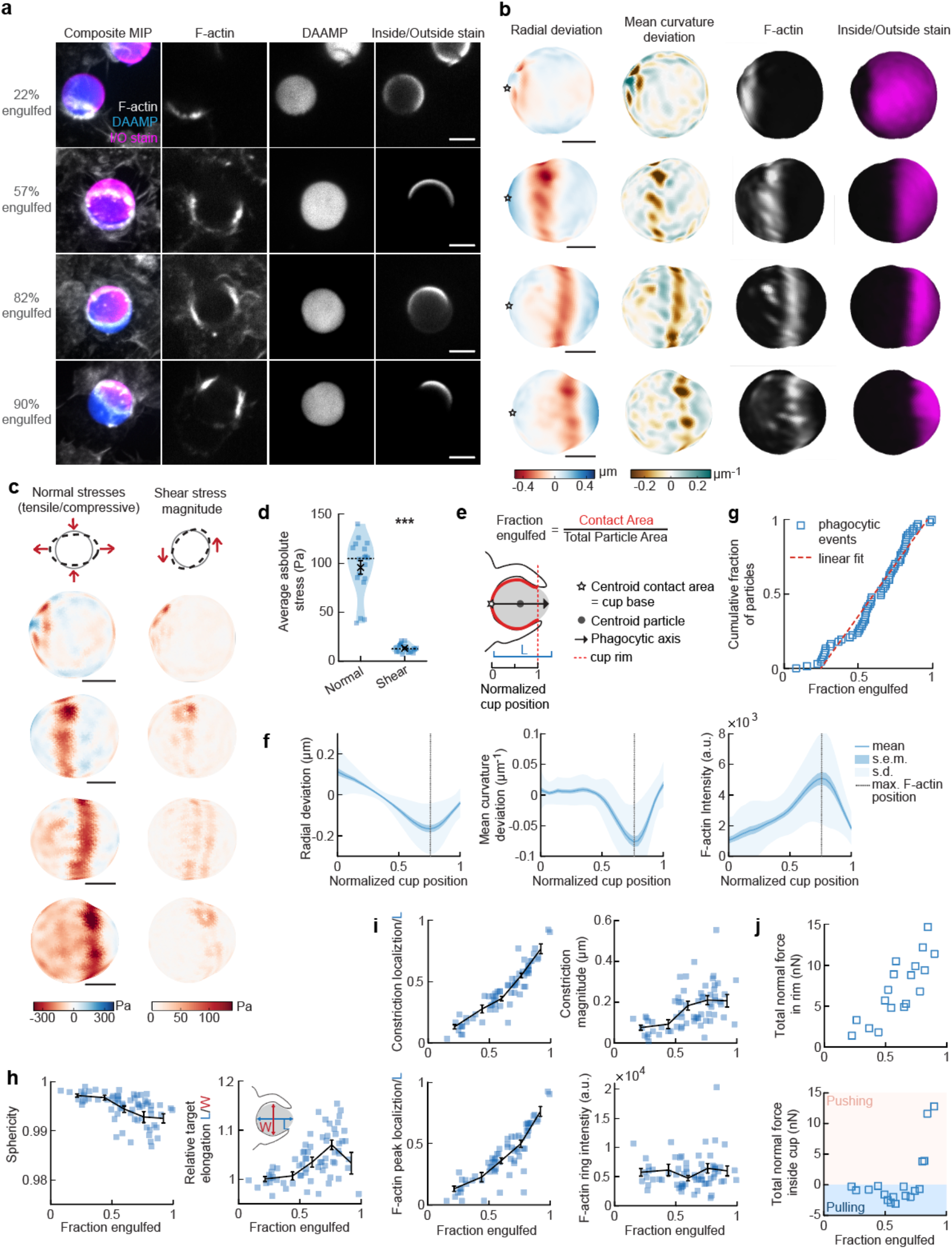
Phagocytic forces include strong actin-mediated constriction and increase with phagocytic progression. **a,** Confocal images of fixed RAW macrophages phagocytosing DAAM-particles functionalized with AF488-Cadaverine, BSA and anti-BSA IgG. Cells were stained for F-actin, and particles with a fluorescent secondary antibody to reveal the exposed surface. Left column: composite maximum intensity projections (MIP) of confocal z-stacks, 2^nd^ to 4^th^ column: single confocal slices through particle centroid. Scale bar, 5 μm. **b,** 3D shape reconstructions of particle in a revealing detailed target deformations and localization of F-actin over the particle surface. Scale bars are 3 μm. Stars mark the base of the phagocytic cup, and the phagocytic axis is horizontal (see **e**). **c,** Normal and shear stresses exerted on the targets in **a,b**. Negative normal forces denote (inward) compressive forces. **d,** Averages of absolute magnitudes of compressive and shear stresses for 18 phagocytic cups spread over various stages of engulfment. Violin plots show individual phagocytic events (blue markers), mean (black cross) and median (dashed line). *Two-side Wilcoxon rank sum test: p = 2.0×10^−4^. **e,** Schematic representation of phagocytic parametrization. Normalized cup position indicates the position along the phagocytic axis relative to the rim of the cup, with 0 the cup base and 1 the rim of the phagocytic cup regardless of engulfment stage. **f,** Average deformation and F-actin intensity profiles along the phagocytic axis to the cup rim. Signals were first processed on a per-particle basis by averaging over the surface along the phagocytic targets in 30 bins. Targets beyond 40% engulfment were included (54 out of 68 events in total). **g,** Cumulative distribution function of the engulfment stage of randomly selected phagocytic events (n = 68). Dashed red line indicates a linear fit. **h,** Target sphericity and elongation depend on phagocytic stage. Blue squares indicate individual measurements, black lines indicate averages within 5 bins. Middle graph inset schematic shows how relative elongation was determined. **i,** Analysis of particle deformation and F-actin fluorescence along the phagocytic axis for all phagocytic events (n = 68). Marker and line styles as in **h**. All error bars indicate s.e.m. unless indicated otherwise. **j,** Analysis of forces in the contractile ring at the cup rim and throughout the remainder of the cup for 18 cups selected for force analysis.

Force analysis revealed compressive stresses up to 400 Pa at these sites, which is substantially greater than our previous findings of stresses of ~ 100 Pa using softer targets with Young’s modulus 0.3 kPa^21^, and suggests that force exertion during phagocytosis may be regulated based on target rigidity. The total compressive forces in the phagocytic rim, which lead to target constriction, increased from ~ 1 nN in early-stage phagocytosis (fraction engulfed ~ 22%) to ~ 10 nN in later stages (fraction engulfed > 50%) (Fig. 2c). The shear forces (Fig. 2c) were consistent in magnitude with reported values using TFM during frustrated phagocytosis on planar gels of similar rigidity^31^ and were ~7-fold lower than the observed normal forces, independent of the stage of phagocytic engulfment (Fig. 2d). This suggests that normal forces dominate the mechanical interaction in phagocytosis, which is in stark contrast with lamellipodial extension during cell migration where shear forces dominate^37,38^.

### Target constriction coincides with the sites of F-actin accumulation and increases with uptake progression

To identify overall trends in F-actin distribution and location of target deformations within the phagocytic cup, we aligned the 3D images of phagocytic cups and analyzed profiles along the phagocytic axis, defined as the axis from the centroid of the cell-target contact area through the target centroid to the opposing target surface (Fig. 2e). This analysis confirmed a clear accumulation of F-actin near the front of the phagocytic cup (~ 5-fold higher than at the cup base), which precisely colocalized with the site of maximal applied inward normal forces regardless of engulfment stage, as illustrated by quantifying the surface mean curvature (Fig. 2f).

To investigate how phagocytic forces change during phagocytic progression, we arranged cups in order by the fraction of their particle surface engulfed, which allowed us to reconstruct phagocytic engulfment over time from fixed cell images (Fig. 2g, Supplementary Fig. 2b). Since DAAMPs cause little optical distortion, measurable features of the phagocytic cups could be analyzed independent of cup orientation and engulfment stage^21^. We found no marked accumulation of cups at any specific stage, suggesting no bottle-neck or rate-limiting steps, which had been previously reported around 50% engulfment^39^. However, we did observe a strong increase in global target deformation, measured as the inverse of target sphericity, with phagocytic progression (Spearman’s ρ = −0.62, p = 5.0 × 10^−8^) (Fig. 2h). This decrease in target sphericity was, at least partially, due to a 4 ± 1% (p = 1.9 × 10^−7^) average increase in DAAMP elongation along the phagocytic axis (Fig. 2h), which is consistent with constriction in the direction orthogonal to the phagocytic axis. Direct analysis of target constriction and F-actin peak intensity for each phagocytic cup (Fig. 2i, Supplementary Fig. 6a) revealed an apparent contractile ring in almost all (~96%) of cups. The location of this actin contractile ring along the phagocytic axis correlated extremely well with phagocytic progression (ρ = 0.93, p = 3.2 × 10^−29^) and led to target constriction increasing from ~80 nm in early-stage (fraction engulfed < 40%) to ~210 nm in late-stage cups (fraction engulfed >70%), which is a direct effect of increasing normal forces at the cup rim (Fig. 2j). Strikingly, in early stages of phagocytosis net pulling (or outward normal) forces were observed throughout the phagocytic cup and particularly at the base (~ 3 nN total force, > 100 Pa tensile stresses) (Fig. 2j), whereas in late-stage phagocytosis strong net compressive stresses were observed.

### Arp2/3-mediated actin polymerization drives force generation throughout phagocytosis, whereas myosin-II powers cup closure

The striking observations of target constriction becoming more pronounced later in engulfment inspired us to consider distinct contributions of actin assembly and myosin-mediated contractility to force generation during phagocytosis. In order to separate the effects of these two actin-dependent processes, we inhibited Arp2/3-mediated and formin-mediated actin polymerization, as well as myosin-II activity using the small molecule inhibitors CK666, SMIFH2 and blebbistatin, respectively (Fig. 3a, Supplementary Fig. 3-5). Target deformation analysis and force calculations revealed that target constriction was strikingly diminished upon inhibition of the Arp2/3 complex and myosin-II activity, while formin inhibition had a relatively modest effect (Fig. 3c-d, Supplementary Fig. 6b, 6d). The loss of target constriction coincided with a strong reduction (~40%) in F-actin accumulation at the rim of the cup, as well as a 50% broadening of the typical narrow (~ 2 μm) F-actin band observed in the DMSO control (Fig. 3b,d,e, Supplementary Fig. 6c). Of note, upon myosin-II inhibition, the loss of F-actin at the rim of the cup was complemented by a small, but significant (p = 0.04), increase in F-actin density at the base of the cup (Fig. 3d,f). This observation suggests that myosin-II may be promoting actin disassembly at the base of the phagocytic cup during internalization, similar to the role of myosin-II in disassembling the F-actin network at the cell rear during cell motility^40^.

**Figure 3:**
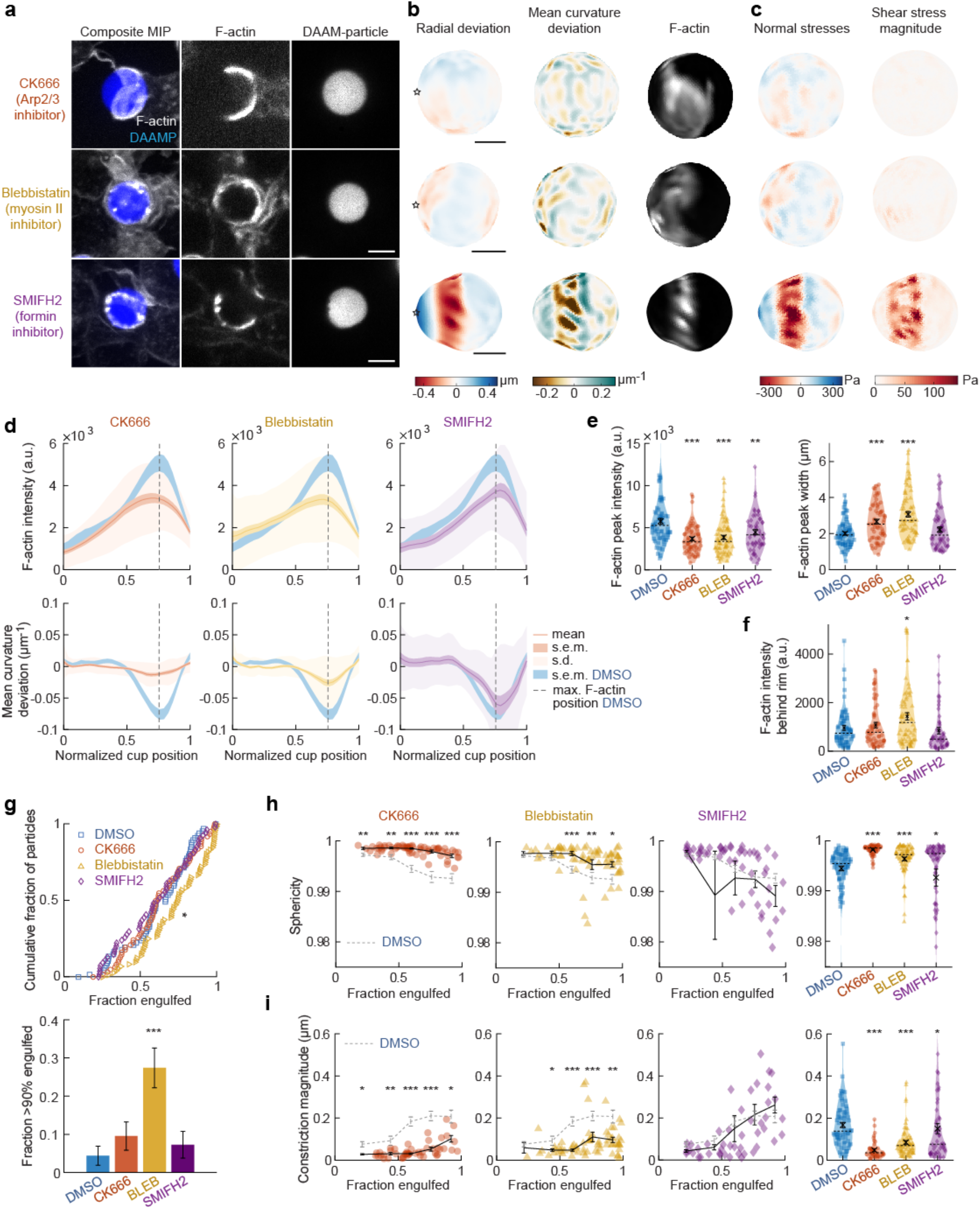
Arp2/3-mediated actin polymerization and myosin-II have distinct roles in phagocytic force generation and progression. **a,** Confocal images of drug-treated fixed RAW cells phagocytosing DAAM-particles functionalized with AF488-Cadaverine, BSA and anti-BSA IgG. Cells were treated with DMSO, CK666 (150 μM), Blebbistatin (15 μM) and SMIFH2 (10 μM) for 30 minutes prior to phagocytic challenge. Each target is approximately 60% engulfed. Fixed cells were stained for F-actin, and particles were labelled with a fluorescent secondary antibody to reveal the exposed surface. Left column: composite maximum intensity projections (MIP) of confocal z-stacks, 2^nd^-3^rd^ column: single confocal slices through particle centroid. Scale bar, 5μm. **b,** Particle shape reconstructions from **a,** revealing detailed target deformations and localization of F-actin over the particle surface. Stars mark the base of the phagocytic cup, and the phagocytic axis is horizontal. Scale bars, 3 μm. **c,** Normal and shear stresses exerted on the target. Negative normal forces denote (inward) compressive forces. **d,** Average deformation and F-actin intensity profiles along the phagocytic axis to the cup rim. Signals were first processed on a per-particle basis by averaging over the surface along the phagocytic targets in 30 bins. Targets before 40% engulfment were excluded. **e, f,** Violin plots of all measured particles, showing individual phagocytic events (colored markers), mean (black cross) and median (dashed line). **e,** F-actin peak intensity and width. **f,** F-actin intensity in the cup (behind the rim), measured right (3μm) behind the main peak for each particle. **g,** Upper panel, cumulative distribution function of the engulfment stage of randomly selected phagocytic events (n = 68, 63, 73 & 55 respectively) from 3 independent experiments. Two sample Kolmogorov-Smirnov test was used (p = 0.016*). Lower panel, fraction late-stage cups. Error bars indicate st.d. estimated by treating phagocytosis as a Bernoulli process. Fisher’s exact test was used to compare fractions (p = 1.9×10^−4^)***. **h,** Sphericity and i, constriction magnitude of DAAM particle changes with phagocytic progression upon drug treatment. Colored markers indicate individual events, black lines indicate averages within 5 bins. Right column, violin plots of all events. Marker and line styles as in **e.** All statistical tests were two-side Wilcoxon rank sum test comparing with the DMSO control (gray) over the same bin with significance levels: p < 0.05*; p < 0.01**; p < 0.001***, unless otherwise indicated. All error bars indicate s.e.m. unless indicated otherwise.

We next investigated whether the activity of these molecular players may be associated with specific phagocytic stages. Our analysis revealed a significant change in the observed distribution of cup stages upon myosin-II inhibition, but not Arp2/3 or formin perturbation, compared to DMSO-treated control cells (Fig. 3g). Specifically, in the blebbistatin-treated cells, we found a > 6-fold enrichment of cups that were beyond 90% engulfment, but not yet closed (p = 1.9×10^−4^). This high prevalence of late-stage phagocytic cups suggests a specific role for myosin-II in cup closure. Throughout phagocytosis, general particle deformations, as measured by the decrease in target sphericity, were strongly reduced upon both CK666 and blebbistatin treatment (Fig. 3h). Inhibition of formins generally reduced target deformations, but also increased the cell-to-cell variability in particle deformation, suggesting that formins may play a role in fine-tuning phagocytic force production (Fig. 3h). Whereas overall target deformations were reduced in all stages of phagocytosis upon Arp2/3 inhibition, myosin-II inhibition only significantly affected phagocytic force generation at later stages, after 50% engulfment (Fig. 3h). A similar effect was observed when quantifying target constriction specifically (Fig. 3i). Thus, in contrast to the prevailing view that myosin-II is dispensable for phagocytosis^7, 27,28^, this analysis strongly suggests that there is a specific role for myosin-II in contributing to the efficiency of late-stage phagocytosis.

### Actin-based protrusive teeth drive target constriction and are mechanosensitive

Based on our observations of discrete spots of inward deformation using MP-TFM (Fig. 1a,b, Fig. 2a,b), and the significant reduction in target deformation after treatment of cells with the Arp2/3 inhibitor CK666, we hypothesized that these local deformations were the result of actin-based protrusions pushing against the surface of the phagocytic target. Indeed, we frequently observed actin rich puncta that appeared as oblong or triangular tooth-like projections locally indenting the target surface along the internal rim of the phagocytic cup (Fig. 4a) and sometimes deeper within the cup (Fig. 4b). Similar actin “teeth” were formed by primary murine bone-marrow derived macrophages (BMDM), bone marrow-derived dendritic cells (BMDC) and HL-60 human neutrophils when challenged with IgG-functionalized DAAMPs, suggesting that these structures are a common feature of phagocytosis (Fig. 4c).

**Figure 4.**
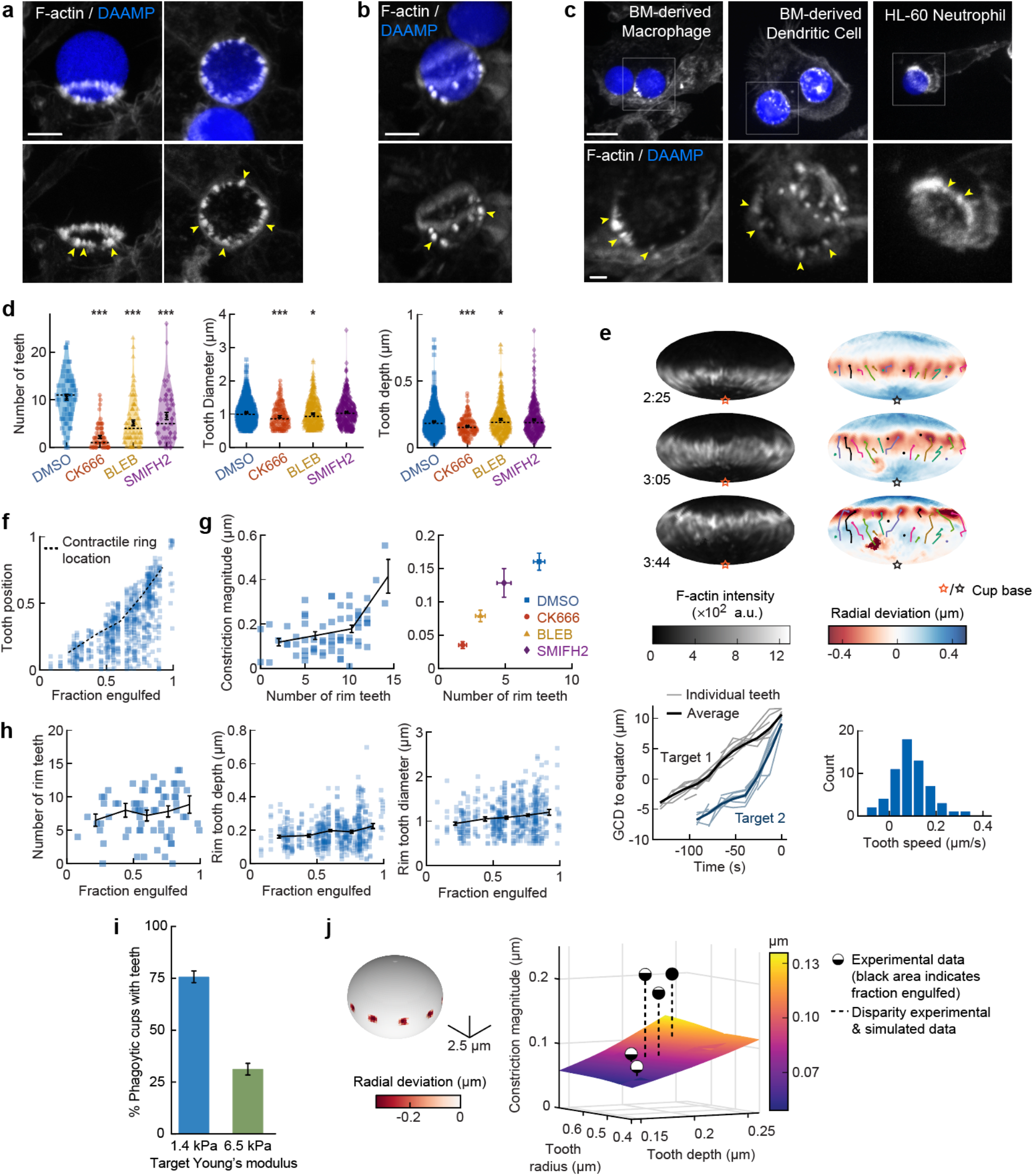
Actin-based teeth are dynamic interconnected structures whose protrusive activity contributes to target constriction. **a,b,** RAW macrophages were fed 1.4 kPa DAAMPs and stained for F-actin. Images represent maximum intensity projections of confocal Z-stacks. Yellow arrows point to actin-based “teeth”. **a,** representative images of teeth at the rim of the phagocytic cup deforming the target. Scale bar, 5 μm. **b,** Example of actin-based teeth observed within the phagocytic cup. Scale bar, 5 μm. **c,** Primary murine bone-marrow derived macrophage (BMDM), as well as primary bone-marrow derived dendritic cell (BMDC) and HL-60 neutrophil-like cells also form actin teeth in response to DAAMP internalization. Scale bar, 5 μm. zoom scale bar, 2 μm. **d,** CK666, blebbistatin, and SMIFH2 treatments reduce formation of actin-based teeth within the phagocytic cup. Teeth size and shape are also modestly affected. Violin plots with individual particles (colored markers), means (black cross), median (dashed line) **e,** Manually tracked actin teeth trajectories from DAAMP internalization imaged by LLSM. Particle surface is shown using Mollweide projection (Supplementary Fig. 2c). Three timepoints of a single phagocytic event are shown, with different colors representing unique teeth. Circles indicate current or final position of a tracked tooth. Lines connect the previous positions of tracked teeth. Lower left; great circle distance (GCD) of teeth to the equator for 2 events. Target one is visualized above, time 0 corresponds to the time at which engulfment was completed. Lower right; distribution of tooth speeds with average 0.094 ± 0.08 μm/s (= 5.6 μm/min) from 3 phagocytic events (60 teeth in total from target 1& 2 in this panel, and from Supplementary Fig. 7d). Tooth speeds were averaged over the trajectory of individual teeth. **f,** Teeth are mostly located at the rim of the cup. Markers represent individual teeth (n = 716) **g,** Constriction magnitude correlates with the number of teeth with Spearman’s rank correlation coefficient (r) = 0.42 (p = 4.4*10^−4^) for individual DMSO phagocytic events (left) and between drug treatments (right). **h,** Teeth number and features change with phagocytic progression, with, from left to right, r = 0.2 (p = 0.11 n.s.), r= 0.17 (p = 1.0×10^−4^), r = 0.16 (p = 2.1×10^−4^). **i,** Phagocytic cups with actin teeth appear more frequently when cells are challenged with softer targets. RAW macrophages were challenged with 9 μm DAAMPs of 1.4 kPa or 6.5 kPa, fixed and stained for F-actin. **j,** Elasticity theory simulations of the relation between tooth size and depth and overall constriction magnitude. Inset shows teeth, simulated as spherical indenters on a spherical target. Bar graph represents pooled data (n =>150 cups) from 3 independent experiments. Error bars indicate st.d. estimated by treating phagocytosis as a Bernoulli process. Pooled data was compared using Fisher’s exact test to compare fractions (p = 1.5×10^−24^***). All error bars indicate s.e.m unless otherwise indicated.

To investigate the nature and biological function of these actin “teeth” more carefully, we identified them on individual particles based on their protrusive nature and high F-actin intensity (Supplementary Fig. 7a,b). According to these criteria, teeth were found in almost all phagocytic cups, with ~10 distinct teeth per cup (Fig. 4d). Typically ~1 μm in diameter, and protruding ~200 nm into the 1.4 kPa DAAMP targets, they resembled podosomes in size and protrusive nature^16^. Cells treated with the Arp2/3 inhibitor CK666, and to a lesser extent cells treated with the formin inhibitor SMIFH2, exhibited a reduction in the number of actin teeth per cup (80% and 40% reduction, respectively) compared to control cells treated with DMSO (Fig. 4d). This result suggests that, like podosomes^17^, target-deforming phagocytic teeth include both Arp2/3- and formin-nucleated actin filaments. Surprisingly, a strong decrease (~50%) in the number of actin teeth was also observed upon myosin-II inhibition. For all treatments, the reduced number of individual teeth that still formed were remarkably similar to those formed by control cells. Although tooth size and depth were reduced upon CK666 or blebbistatin treatment, the effect size was small (< 15%), suggesting that “teeth” are resilient structures that, once formed, have well-defined properties (Fig. 4d).

LLSM allowed us to track the dynamics of actin teeth during phagocytosis (Fig. 4e), which revealed clear forward movement over the target surface. A few teeth that we initially detected near the rim stayed in place, remaining where they had assembled during early-stage phagocytosis, suggesting that teeth located deeper within the phagocytic cup (as observed in the fixed cell images) may have originated earlier from the cup rim and been left behind as the cup progressed (Fig. 4b, Supplementary Fig. 7e). More commonly, however, teeth moved forward with a speed of ~5.6 μm/min, similar to the previously reported values for podosome-like structures on very stiff substrates during frustrated phagocytosis^41^. Strikingly, teeth within the same phagocytic cup appeared to move in a coordinated fashion, with similar speed and direction, and even with observed collective speed changes (Fig. 4e, Supplementary Fig. 7d). This suggests that phagocytic teeth, like podosomes^42,43^, are mechanically interlinked at the mesoscale.

To test whether the actin teeth were mechanosensitive, we challenged RAW macrophages to ingest 9 μm DAAMPs of 1.4 or 6.5 kPa and fixed and stained cells to examine actin teeth formation. Interestingly, RAW macrophages assembled actin teeth more frequently when fed softer targets (Fig. 4i), suggesting that phagocytic teeth may play a role in the overall mechanosensitivity of phagocytosis^8,18–21^.

Given the ring-like organization of phagocytic teeth in the cup rim, combined with their individual protrusive activity, we questioned whether they were sufficient to explain our observations of target constriction orthogonal to the phagocytic axis, or if a separate contractile mechanism is required. We first distinguished the teeth positioned at the rim of the cup (~70% of teeth), which likely contribute to target constriction, from those deeper in the cup, based on their distance from the cup rim (Fig. 4f, Supplementary Fig. 7c). We then determined whether the properties of teeth near the rim correlated with the overall target constriction. Indeed, the number of teeth per cup and tooth size correlated with overall constriction in DMSO treated cells and between groups treated with actomyosin activity inhibitors CK666, SMIFH2 and Blebbistatin (Fig. 4g, Supplementary Fig. 7f). We further examined whether changes in the teeth could be related to increasing target constriction with phagocytic cup progression. Teeth numbers increased only slightly with phagocytic progression, which suggests that they are formed quickly early on in phagocytosis and are then typically maintained at constant numbers throughout engulfment (Fig. 4h). Teeth size and depth increased significantly but modestly during phagocytic progression (Fig. 4h, Supplementary Fig. 7g). Elasticity theory simulations of teeth-like indentations of a spherical target allowed us to test whether teeth protrusive activity is sufficient to explain the extent of overall target constriction in different stages of phagocytosis (Supplementary Fig. 8). Remarkably, this revealed that teeth activity is indeed sufficient to account for total target constriction in early-stage phagocytosis (< 50% engulfment), but insufficient to explain the greater degree of target constriction later in the process (Fig. 4j). This is consistent with additional myosin-II based contractile forces in late-stage phagocytosis, as suggested by our observations using Blebbistatin.

### Regulators of branched actin assembly (Arp2/3 and WASP) and myosin-I isoforms localize to actin teeth while myosin-II forms contractile rings within the phagocytic cup

Due to the resemblance of the phagocytic actin teeth to podosomes in size, protrusive activity (Fig. 4d) and dynamics (Fig. 4e), we naturally questioned whether these structures were similar in protein composition as well. Given technical challenges with immunohistochemical staining and MP-TFM (see methods), we transfected the RAW macrophages with fluorescently tagged proteins to assess tagged protein localization relative to the actin teeth using 3D reconstructions of the DAAMP (Fig. 5a-b). Consistent with our earlier results showing a decrease in the number of teeth after treating cells with the Arp2/3 inhibitor, we found that the Arp2/3 complex and its activator, WASP, colocalized with the actin teeth. Moreover, cortactin and cofilin, actin-binding proteins that are frequently found in association with densely branched actin networks, also localized to the phagocytic teeth (Supplementary Fig. 9a). In contrast, myosin-II often appeared distinctly behind the actin teeth in an anti-correlated fashion (Fig. 5b). In particular, rings composed of myosin-II filaments could be seen within the phagocytic cup (Fig. 5a). In phagocytic cups that were further along the engulfment process, myosin-II coalesced into a ring behind the actin rim (Supplementary Fig. 9b) and clearly localized at sites of target constriction in cases of extreme target deformation (Fig. 5c). Meanwhile, the two long-tailed myosin-I isoforms, myo1e and myo1f, localized specifically to the tips of the actin teeth, consistent with our previous observations (Fig. 5a)^12^. Two formins tested, mDia2 and FHOD1, did not localize to phagocytic actin teeth (Supplementary Fig. 9c).

**Figure 5.**
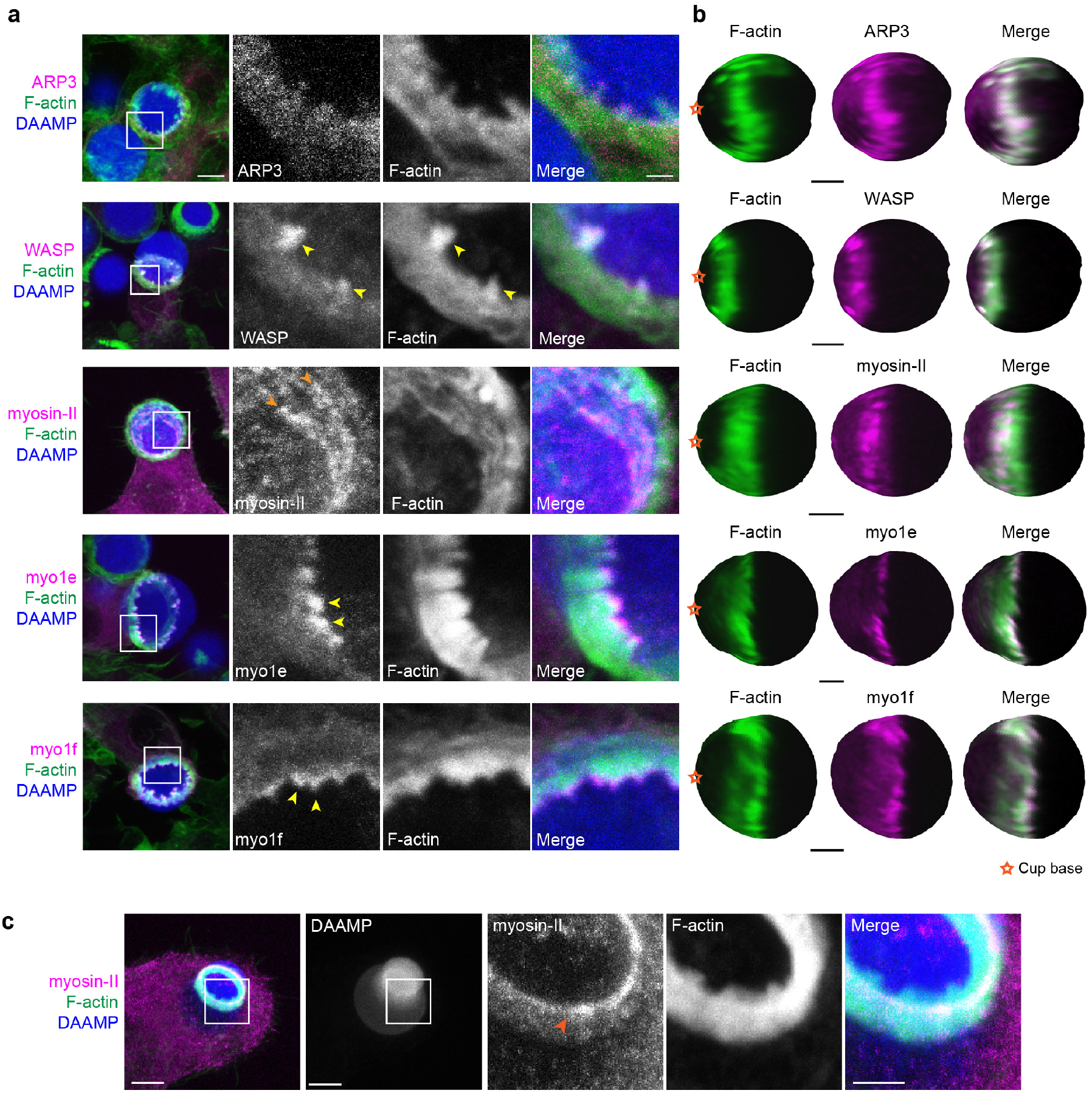
Multiple actin regulatory proteins localize to phagocytic teeth. **a,** RAW macrophages were transfected with fluorescently tagged actin binding proteins and challenged to ingest DAAMPs (11 μm, 1.4 kPa) functionalized with AF647-Cadaverine, BSA and anti-BSA IgG to assess localization to actin teeth (yellow arrowheads). Images are maximum intensity projections of confocal Z-stacks. White boxes in leftmost panels indicate the site of the zoomed images to the right. Scale bar, 5 μm. Zoom scale bar, 1 μm. **b,** DAAM-particle reconstructions for examples shown in **a,** showing target deformations and localization of fluorescent proteins with respect to actin teeth. Scale bar, 3 μm. **c,** Myosin-II condensing into thick concentric rings (marked by orange arrowheads) during late-stage phagocytosis of a highly deformed target. Images are maximum intensity projections of confocal Z-stacks. Scale bar, 5 μm. Zoom scale bar, 1 μm.

Given both recent and older observations identifying adhesion adaptor proteins at the phagocytic cup^18,44^, we were particularly interested in the localization of paxillin and vinculin. In comparison to the branched actin-binding proteins, both paxillin and vinculin localized behind the phagocytic teeth in a punctate-like pattern (Supplementary Fig. 9d). Altogether, these studies support a model where the actin teeth are composed of branched actin filaments guided by myosin-I motor proteins. The localization of myosin-II and paxillin/vinculin behind the teeth suggests that these proteins may contribute to the mechanical interconnection of the teeth, similar to podosomes on 2D surfaces^17,43^.

### Contractile activity may enable resolution of phagocytic conflicts via partial target eating (“nibbling”) or forfeit of uptake (“popping”)

While we have found that target constriction is a signature mechanical feature of FcR-mediated phagocytic progression, it is unclear what functional role target constriction might play during phagocytosis, since actin-driven membrane advancement along the target surface should in principle be sufficient for internalization^45,46^. In addition to the many successful internalization events we observed using LLSM, we also observed some strikingly different target encounters in which RAW macrophages assembled large amounts of F-actin, only to squeeze futilely at the base of the target without completing engulfment. In these cases, the contractile activity resulted in dramatic deformations and even dumbbell-like appearance of the target (Fig. 6a,b, Supplementary Video 5). In addition to this kind of internalization failure by single cells, we also observed incidents where two macrophages engaged one DAAMP target. Similar to previous observations of red blood cells (RBCs) being squeezed into multilobed shapes when attacked by two macrophages simultaneously^22^, these conflicts were also observed using primary BMDMs challenged with DAAMPs (Supplementary Video 6). Although the polymeric targets used in this study prohibit partial target eating because each particle is effectively one single crosslinked macromolecule that cannot easily be severed by cell-scale forces, this behavior may be reminiscent of trogocytosis, the process which has been observed during immune cell attack of cancer cells whereby phagocytes ingest small bits of their target^47–49^. By imaging RAW macrophages transfected with GFP-NMMIIA, we observed highly enriched rings of myosin-II signal at DAAMP deformations during attempts of partial target eating (Fig. 6c, Supplementary Video 7). Target encounters involving extreme deformations of the DAAMP also revealed the existence of a “popping” mechanism that could lead to a sudden release of the target (Supplementary Video 8) or, conversely, a sudden completion of engulfment (Supplementary Video 9, Supplementary Fig. 10). During such events, targets were first gradually deformed to a dumbbell-like shape, followed by a sudden translocation of the particle, as well as an immediate recovery of its original spherical shape and volume (Fig. 6d, Supplementary Video 7). The rapid timescale of this process suggests that it is likely purely mechanical, representing an elastic recoil of the DAAMP. Importantly, these encounters were rather common, with the attempted partial eating attempts (~14%) and popping (~24%) making up almost 40% off all recorded events (Fig. 6e). Furthermore, such events, and specifically popping, were mechanosensitive and occurred much less frequently for stiffer 6.5 kPa targets (~1%, n = 89, p = 1.5 × 10^−6^), resulting in the overall more frequent failure of phagocytosis for soft particles (Fig. 6f).

**Figure 6.**
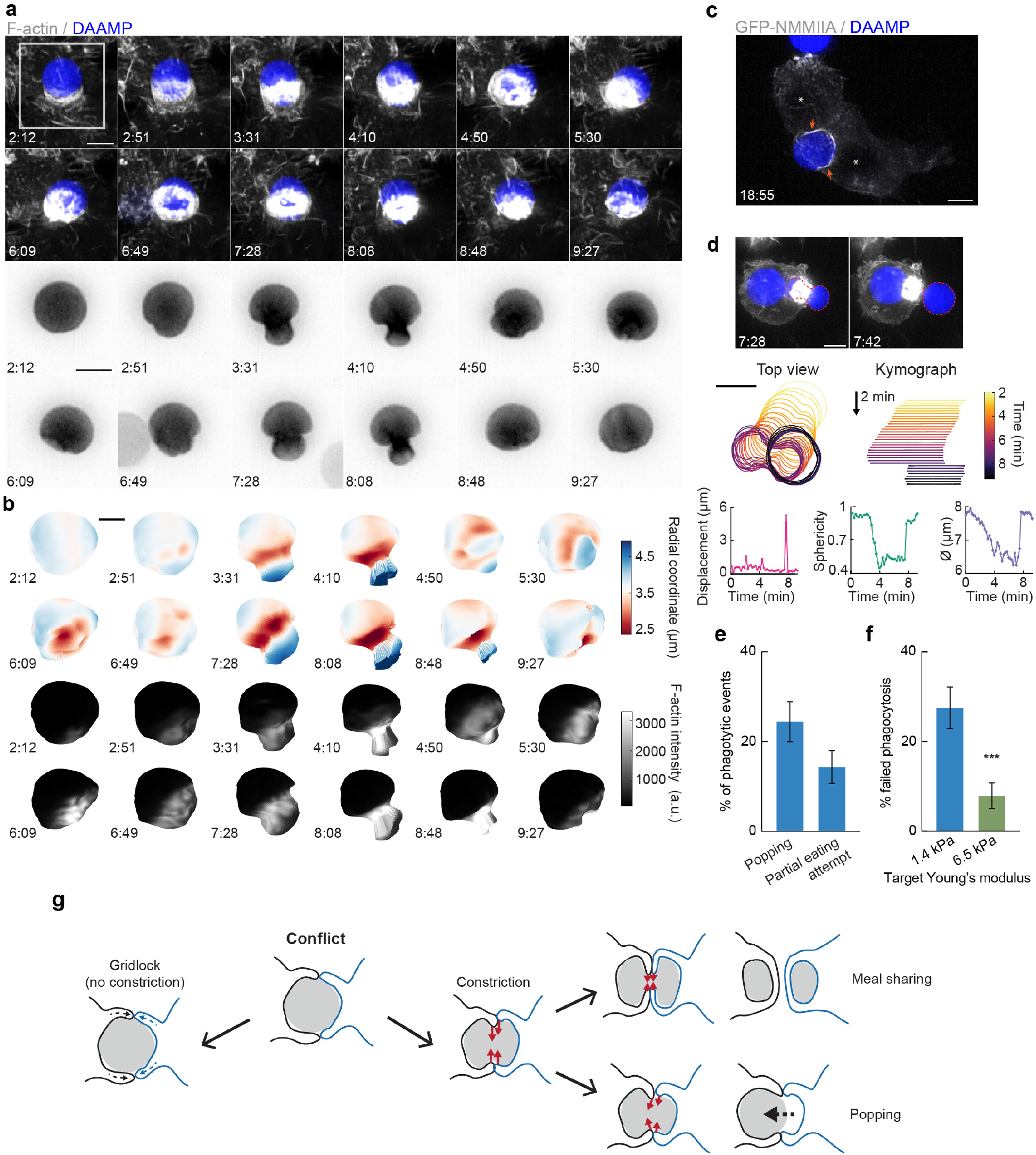
Contractile activity may enable resolving conflicts by partial target eating and a popping release mechanism. **a,** Maximum intensity projections (MIP) of LLSM stacks showing failed internalization attempt of RAW macrophage with IgG-functionalized 1.4 kPa DAAMP. Zoomed images of the area marked by the white box showing only the DAAMP channel (inverse grayscale LUT) are shown below. **b,** 3D reconstructions showing both the particle shape and F-actin signal over the particle surface corresponding to the event in **a.** Scale bar, 3 μm. **c,** MIP of LLSM stacks showing myosin-II accumulation (orange arrows) during a partial eating event. RAW macrophages (marked by *) were transfected with EGFP-NMMIIA and challenged with IgG-functionalized 1.4 kPa DAAMP. **d,** Top: MIPs of LLSM stacks of RAW macrophage suddenly releasing heavily deformed target. Red dashed line outlines DAAMP. Middle, particle position and outline (left), and kymograph of particle position (right). Bottom, Particle displacement, sphericity and apparent diameter over time of the same event shows the sudden nature of the release. **e,** Sudden forfeit by a popping mechanism and attempted partial eating are common for 1.4 kPa targets, with ~24% and ~14% occurrence of all phagocytic events, respectively. **f,** Percentage of failed phagocytic events is dependent on particle rigidity. Data from 2-4 independent experiments was pooled (n = 89, 91 phagocytic events) and compared using Fisher’s exact test (p = 7.4×10^−4^). **g,** Schematic representation of the multiple ways in which target constriction may enable resolving macrophage conflicts in which two cells attempt a single target. All scale bars are 5 μm, unless otherwise indicated. All error bars indicate s.d. estimated by treating phagocytosis as a Bernoulli process.

## Discussion

Through the combination of LLSM and MP-TFM, we report here a detailed analysis of the mechanical progression of phagocytosis and the contributions of several key molecular players. Most importantly, we have discovered that FcR-mediated phagocytosis occurs through a unique mechanism in which normal forces dominate over shear forces in the cell-target interaction. This is in contrast to the current view that the phagocytic cup is equivalent to the leading edge of a migrating cell, where shear forces typically predominate at the cell-substrate interface^37,38^. In addition, the fast forward movement of actin teeth, which underlie target constriction, concomitant with phagocytic cup progression across the target is in stark contrast to lamellipodial focal adhesion complexes (FAs), which are fixed relative to the substratum^38^. The strong target constriction recently observed in complement-mediated phagocytosis by peritoneal macrophages *in vitro*^50^, and in phosphatidylserine-mediated phagocytosis by epithelial cells in zebrafish embryos^51^ suggests that strong normal forces and target constriction are likely a general feature of phagocytosis.

We show that normal forces are primarily generated by protrusive phagocytic actin “teeth” and myosin-II contractility, which make distinct contributions to target constriction. While actin-based puncta have been previously observed during phagocytosis of stiffer polystyrene beads^12,13^, their presence in multiple phagocytic cell types (Fig. 4c) suggests a common role in phagocytosis. These structures are podosome-like in protein composition, consisting of mostly branched F-actin and actin regulatory proteins. Accordingly, inhibition of the Arp2/3 complex dramatically reduces tooth number and size and consequently reduces target constriction. Unexpectedly, myosin-II motor inhibition also reduces tooth number and size, albeit to a lesser extent than Arp2/3 inhibition, and also alters tooth spatial distribution (Supplementary Fig. 7c). In podosomes, myosin-II filaments interconnect radial actin fibers of individual podosomes to create a coherent network^43^. The localization of myosin-II behind and in between teeth, as well as the coordinated movement of teeth, suggests that actin teeth in phagocytosis are similarly interconnected by actomyosin structures. This molecular arrangement of actin teeth (Fig. 7a) bears similarity to the organization of podosome rosettes^14^, and we show that they possess a similar mechanosensitivity (Fig. 4j)^15–17^. We find that individual teeth grow larger and stronger with phagocytic progression, correlating with increasing myosin-II constriction observed in late-stage phagocytosis (Fig. 7a). These observations may relate to the force feedback mechanism observed in Arp2/3 mediated branched actin networks *in vitro* showing that mechanical resistance makes self-assembling actin networks stronger^52^. Thus, increased myosin-II contraction in late-stage phagocytosis may promote stronger actin teeth. This rapid structural reinforcement by myosin-II has not been described for actin cores of podosomes on 2D, making this feature unique to phagocytosis.

**Figure 7.**
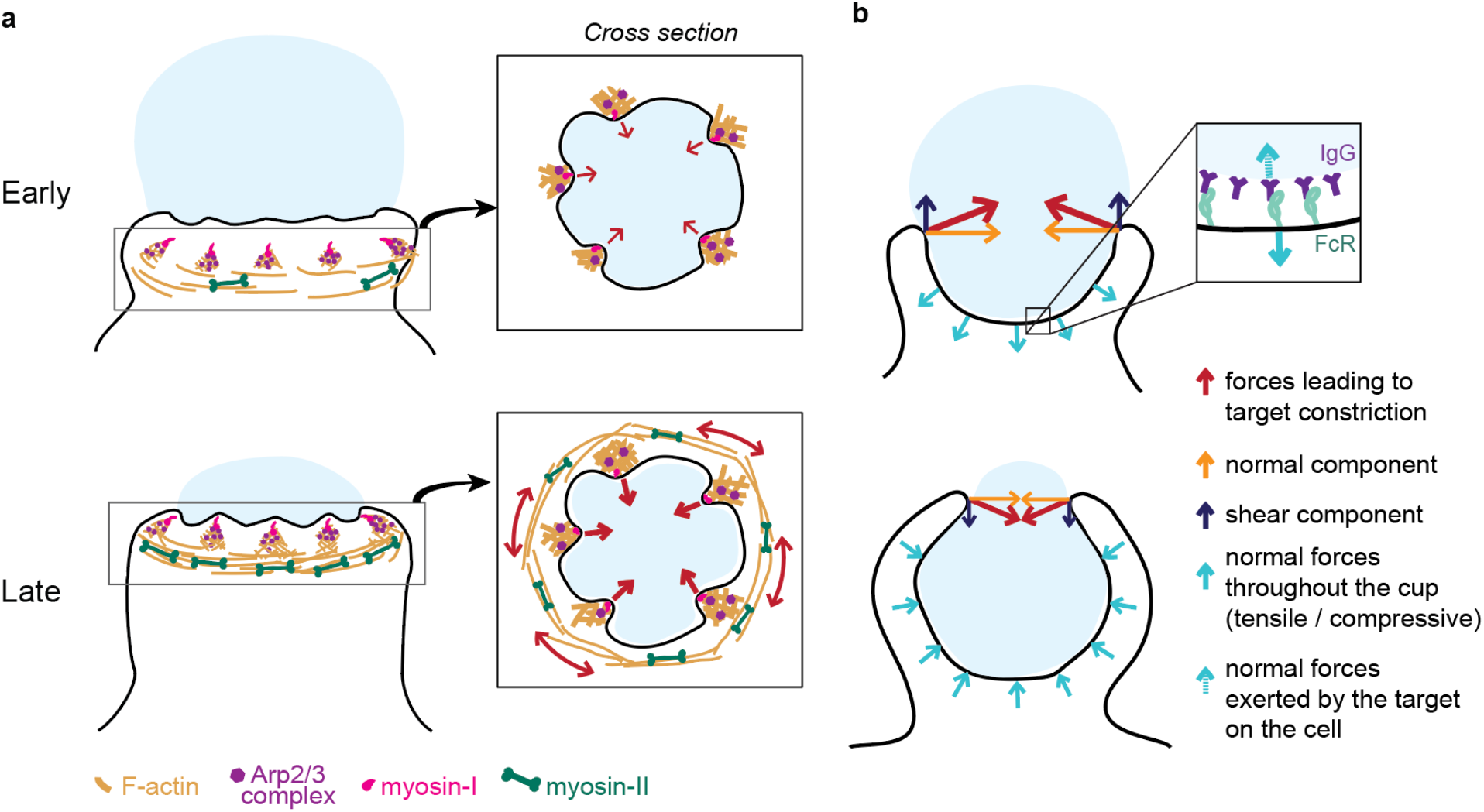
Model of the molecular players and force balance during phagocytosis. **a,** Graphical model: Arp2/3 mediated actin teeth, guided by myosin-I motors, and organized in a ring drive phagocytic progression and inward target deformation. Teeth are interconnected through myosin-II filaments located behind the actin teeth. **b,** Ring-like target constriction by protrusive actin teeth and myosin II activity (red arrows) can be decomposed in forces orthogonal to the phagocytic axis, which are balanced within the ring (yellow arrows) and along the phagocytic axis, which result in a net force exerted by the ring (purple arrows). The local target geometry at the protruding rim of the cup determines the direction of the net force. Before 50% engulfment, this force points outward and is balanced by pulling forces throughout the base of the phagocytic cup (push and lock). Inset, the pulling forces from the cell (dashed arrow) on the target and paired forces from the target on the cell (solid arrow) likely put the receptor-ligand interactions under tension. After 50% engulfment, the net ring force points inward, and is balanced by compressive forces (cyan arrows) throughout the phagocytic cup.

We further show that myosin-II plays an important role in phagocytic cup closure – a stage in phagocytosis that has been notoriously difficult to study experimentally^53^ (Fig. 7a). Generally, myosin-II is not deemed important for cup closure or phagocytic progression^7,22^, yet we observe myosin-II enrichment at the rim of phagocytic cups (Fig. 5c). Moreover, blebbistatin treatment specifically affects late-stage contractile force generation (Fig. 3h) causing cup closure to become a bottle-neck step as evidenced by the accumulation of late-stage phagocytic cups (Fig. 3g). Because these forces are exerted normal to the target surface, the contribution of myosin-II activity to phagocytic progression has likely gone unnoticed in previous studies using regular TFM on a planar substrate, which only measures forces that are tangential to the surface (shear)^12,20,25^.

We show that phagocytosis is hallmarked by a unique force balance, where before 50% engulfment, target constriction results in an outward force balanced by pulling forces throughout the base of the phagocytic cup (Fig. 7b). Given the reduced presence of cytoskeletal components at the base of the cup (Supplementary Fig. 2b)^4,7^, these pulling forces are most likely not due to actin polymerization forces or actomyosin contractility, but instead a result of the target being held in place through receptor-ligand bonds throughout the base of the cup. Hence, the forces acting on receptor-ligand bonds within the cup are likely dependent on the target physical properties and the local geometry at the rim of the cup (Fig. 7b). Since the lifetime of such bonds is tension-dependent^54,55^, this push-and-lock mechanism may enable sensing of physical target properties through a proofreading mechanism and thereby aid macrophages in target selection. Indeed, we observe that forfeit of uptake via popping, in which the outward-directed forces likely overcome the strength of the receptor-ligand interactions, depends on the local target geometry and target rigidity. In addition, for non-spherical targets (*e.g.* ellipsoidal), this force balance would result in a net torque aligning the target’s long axis along the phagocytic axis, which has been observed previously and is reported to lead to enhanced uptake efficiency of ellipsoidal targets^19,56,57^.

We observed that actin teeth formed more frequently when cells were challenged with softer targets, yet we also associated softer targets with more instances of failed internalization. This calls into question the effectiveness of actin teeth, and, more generally, target constriction, in driving phagocytic internalization. Aside from a role in cup closure, target constriction could be important for creating a tight apposition between the cell and target, which is essential for receptor engagement^58^. Surprisingly though, we noticed that overly strong target constriction leads to failure of attempted phagocytosis, expressed either as partial eating or a popping mechanism leading in forfeit of uptake. Although these mechanisms may lead to reduced uptake efficiency in isolated phagocyte-target interaction, we suspect they may be critical in more complex phagocytic encounters that occur *in vivo*, *e.g.* when multiple macrophages approach a single target (Fig. 6g), or attempt to engulf adherent and hard-to-reach targets^8,11,59^. Altogether, our findings show that actin polymerization-dependent protrusive forces and myosin-II-dependent contractile forces contribute to driving target deformation and phagocytic internalization, and likely both participate in the mechanosensation required for phagocytic plasticity.

## Code availability

The Matlab code for analysing confocal images and deriving particle shape is publicly available on https://gitlab.com/dvorselen/DAAMparticle_Shape_Analysis. The Python code used for analysing tractions is provided on https://gitlab.com/micronano_public/ShElastic.

## Acknowledgements

This work was supported by American Heart Association (18PRE34070066) grant to S.R.B., the National Institute of Diabetes and Digestive and Kidney Diseases of the NIH under Award R01DK083345 to M.K., the Italian Association for Cancer Research (AIRC), Investigator Grant (IG) 20716 to N.C.G., and the Howard Hughes Medical Institute (J.A.T.). D.V. further acknowledges the Cancer Research Institute for support through a CRI Irvington fellowship. LLSM imaging was performed at the Advanced Imaging Center (AIC)—Howard Hughes Medical Institute (HHMI) Janelia Research Campus. We thank John M. Heddleston and Jesse Aaron of the AIC for helpful discussion. The AIC is jointly funded by the Gordon and Betty Moore Foundation and the Howard Hughes Medical Institute. We thank Lorenzo L. Labitigan for critical review of the manuscript and Sharon Chase for help with animal experiments.

## Author Contributions

S.R.B., D.V., N.C.G., J.A.T. and M.K. conceived of and designed the study. S.R.B performed experiments and D.V. generated reagents, wrote the image analysis software and analyzed the data. Y.W. and W.C. performed the force analysis. J.A.T., N.C.G. and M.K. supervised the project. S.R.B., D.V., N.C.G., J.A.T., and M.K. wrote the manuscript.

## Methods

### Cell culture

RAW264.7 (ATCC; male murine cells) were cultured in DMEM, high glucose, containing 10% FBS and 1% antibiotic-antimycotic (Gibco) (cDMEM) at 37 °C with 5% CO_2_. HL-60 cells (ATCC; CCL-240) were cultured in RPMI plus L-glutamine supplemented with 20% FBS and 1% antibiotic-antimycotic (cRPMI). HL-60 cells were differentiated into neutrophil-like cells in culture media containing 1.5% DMSO and used at day 5-6 post-differentiation and plated on 20 μg/mL fibronectin. For the collection of primary murine bone marrow progenitor cells, femurs and tibias of C57BL/6 mice were removed and flushed with cDMEM. Red blood cells were lysed using ACK buffer (0.15 M NH_4_Cl) and bone marrow progenitor cells were recovered by centrifugation (250×g, 5 min, 4 °C), washed once with sterile PBS and plated on tissue culture dishes in cDMEM at 37 °C with 5% CO_2_. For differentiation into bone-marrow derived macrophages (BMDM), non-adherent cells were moved to bacteriological Petri dishes the next day and differentiated over 1 week in cDMEM containing 20 ng/mL recombinant murine M-CSF (Biolegend, 576404). Generation of murine bone-marrow derived dendritic cells (BMDC) has been previously described^60^. In brief, bone marrow progenitor cells were collected in cRPMI and replated in cRPMI containing 20 ng/mL recombinant murine GM-CSF (Peprotech, 315-03) (DC media). On day 3, DC media was supplemented. On days 6 and 8, half of the culture supernatant and nonadherent cells were spun down and resuspended in cRMPI containing 5 ng/mL GM-CSF. DC maturation was assessed on day 10 by flow cytometry using PE-Cd11c (Biolegend, 117307) and FITC-MHC-II (Biolegend, 107605) staining. Cells were used on Day 12. All procedures utilizing mice were performed according to animal protocols approved by the IACUC of SUNY Upstate Medical University and in compliance with all applicable ethical regulations.

### Chemicals and drugs

Blebbistatin, CK-666, and SMIFH2, and fibronectin were purchased from EMD Millipore. Alexa Fluor-488, Alexa Fluor-568, Alexa Fluor-647 conjugated phalloidin were purchased from Life Technologies. Janelia Fluor 549 (JF549) HaloTag Ligand was a generous gift from Luke Lavis.

### Constructs and transfection

Human myo1e and myo1f constructs tagged with EGFP, mEmerald-C1, or mScarlet-C1 have been previously described^12^. mEmerald-Lifeact was a gift from Michael Davidson (Addgene #54148). Chicken regulatory light chain (RLC) tagged with EGFP was a gift from Klaus Hahn. WASP tagged with myc was a gift from Dianne Cox, and was subcloned into pUB-Halo-C1 vector. CMV-GFP-NMHCII-A was a gift from Robert Adelstein (Addgene #11347). ARP3-mCherry (Addgene #27682) and mCherry-cortactin (Addgene #27676) were gifts from Christien Merrifield that were subcloned into EGFP-C1 and mEmerald-C1, respectively. Chicken paxillin was a gift from Chris Turner, which was subcloned into mScarlet-i-C1. Chicken vinculin was a gift from Kenneth Yamada (Addgene #50513) and subcloned into pUB-mEmerald-C1. Immunohistochemical staining to determine localization of select podosome-related proteins in relation to the actin teeth did not produce good results, which may be due to the adhesive and porous nature of the IgG-functionalized DAAMPs. As an alternative, we transfected the RAW macrophages with fluorescently tagged proteins. All transfections were accomplished by electroporation (Neon) using the manufacturer’s instructions.

### Microparticle synthesis

DAAM-particles were synthesized as previously described^21^. First, acrylamide mixtures containing 100 mg/mL acrylic components, 150 mM NaOH, 0.3% (v/v) tetramethylethylenediamine (TEMED), 150 mM MOPS (prepared from MOPS sodium salt, pH 7.4) were prepared. Mass fraction of acrylic acid was 10% and crosslinker mass fraction was 0.65% or 2.3%, for 1.4 kPa and 6.5 kPa particles, respectively. Prior to extrusion, the gel mixture was degassed for 15 minutes and then kept under nitrogen atmosphere until the extrusion process was complete. Tubular hydrophobic Shirasu porous glass (SPG) were sonicated under vacuum in n-heptane, mounted on an internal pressure micro kit extruder (SPG Technology Co.) and immersed into the oil phase (~125 mL) consisting of hexanes (99%) and 3% (v/v) Span 80 (Fluka, 85548 or Sigma-Aldrich, S6760). 10 mL of gel mixture was extruded through SPG membranes under nitrogen pressure of approximately 7 kPa, 15 kPa, for membranes with pore size 1.9 μm and 1.4 μm, respectively. 9 mm, 1.4 kPa particles were synthesized using 1.4 μm pore size membranes, whereas 9 μm, 6.5 kPa particles and 11 μm, 1.4 kPa particles were made using 1.9 μm pore size membranes. The oil phase was continuously stirred at 300 rpm and kept under nitrogen atmosphere. After completion of extrusion, the emulsion temperature was increased to 60 °C and polymerization was induced by addition of ~225 mg 2,2’-Azobisisobutyronitrile (AIBN) (1.5 mg/mL final concentration). The polymerization reaction was continued for 3 h at 60 °C and then at 40 °C overnight. Polymerized particles were subsequently washed (5× in hexanes, 1× in ethanol), dried under nitrogen flow for approximately 30 minutes, and resuspended in PBS (137 NaCl, 2.7 mM KCl, 8.0 mM Na_2_HPO_4_, 1.47 mM KH_2_PO_4_, pH 7.4) and stored at 4 °C.

### Microparticle functionalization

DAAM-particles were functionalized as previously described^21^. In brief, DAAMPs were diluted to 5% (v/v) concentration and washed twice in activation buffer (100 mM MES, 200 mM NaCl, pH 6.0). They were then incubated for 15 min in activation buffer supplemented with 40 mg/mL 1-ethyl-3-(3-dimethylaminopropyl) carbodiimide, 20 mg/mL N-hydroxysuccinimide (NHS) and 0.1% (v/v) Tween 20, while rotating. Afterwards they were spun down (16,000 × g, 2 min) and washed 4X in PBS, pH 8 (adjusted with NaOH) with 0.1% Tween 20. Immediately after the final wash the particles were resuspended in PBS, pH 8 with BSA (Sigma, A3059) and incubated, rocking for 1h. Then cadaverine-conjugate was added: either Alexa Fluor 488 Cadaverine (Thermo Fischer Scientific, A-30679) or Alexa Fluor 647 Cadaverine (Thermo Fischer Scientific, A-30676) to a final concentration of 0.2 mM. After 30 min, unreacted NHS groups were blocked with 100 mM TRIS; 100 mM ethanolamine (pH 9). DAAM-particles were then spun down (16,000 × g, 2 min) and washed 4X in PBS, pH 7.4 with 0.1% Tween 20. Finally, BSA-functionalized DAAMPs were resuspended in PBS, pH 7.4 without Tween.

### Phagocytosis assay

DAAM particles were washed 3X in sterile PBS and opsonized with rabbit anti-BSA antibody (MP Biomedicals, 0865111) for 1 h at room temperature. DAAMPs were then washed three times (16000 x g, 2 min) with PBS and resuspended in sterile PBS. DAAM particles were added to a total volume of 400 μl of serum-free DMEM, briefly sonicated in a bath sonicator, and applied to phagocytes in a 12-well plate. To synchronize phagocytosis and initiate DAAMP-phagocyte contact, the plate was spun at 300 × *g* for 3 min at 4°C. Cells were incubated at 37°C to initiate phagocytosis for a period of 3-5 min. Media was then removed and cells were fixed with 4% PFA/PBS for 15 min. Any unbound DAAMPs were then washed away with 3X washes of PBS and cells were stained with goat anti-rabbit-Alexa Fluor-405 antibodies (Invitrogen, A31556, 1:400) for 30 min to visualize exposed DAAM area. Cells were then washed with PBS (3 × 5 min) and permeabilized with 0.1% Triton X-100/PBS for 3 min, then stained with Alexa Fluor-568 or −488 conjugated phalloidin (1:300). Coverslips were then mounted using VECTASHIELD Antifade Mounting Medium (Vector Laboratories, H-1000) and sealed with nail polish. For drug treatments, cells were exposed to the indicated drug concentration for 30 min prior to the assay and DAAM particles were resuspended and exposed to cells in the same drugged media.

### Microscopy

Confocal images were taken using a PerkinElmer UltraView VoX Spinning Disc Confocal system mounted on a Nikon Eclipse Ti-E microscope equipped with a Hamamatsu C9100-50 EMCCD camera, a 100 × (1.4 N.A.) PlanApo objective, and controlled by Volocity software. Images for protein localization were taken using a Leica TCS SP8 laser scanning confocal microscope with a HC Pl APO 63×/1.4 NA oil CS2 objective at Upstate/Leica Center of Excellence for Advanced Light Microscopy. Confocal image data were less prone to artifacts than the LLSM images (Supplementary Fig. 1j), and therefore chosen for accurate force analysis.

The lattice light-sheet microscope^61^ utilized was developed by E. Betzig and operated/maintained in the Advanced Imaging Center at the Howard Hughes Medical Institute Janelia Research Campus (Ashburn, VA). 488, 560, or 642 nm diode lasers (MPB Communications) were operated between 40 and 60 mW initial power, with 20–50% acousto-optic tunable filter (AOTF) transmittance. The microscope was equipped with a Special Optics 0.65 NA/3.75 mm water dipping lens, excitation objective and a Nikon CFI Apo LWD 25 × 1.1 NA water dipping collection objective, which used a 500 mm focal length tube lens. Live cells were imaged in a 37 °C-heated, water-coupled bath in FluoroBrite medium (Thermo Scientific) with 0–5% FBS and Pen/Strep. Images were acquired with a Hamamatsu Orca Flash 4.0 V2 sCMOS cameras in custom-written LabView Software. Post-image deskewing and deconvolution was performed using HHMI Janelia custom software and 10 iterations of the Richardson-Lucy algorithm.

### Microparticle 3D shape reconstruction and force analysis

Image analysis was performed with custom software in Matlab, similar to described previously^21^. Briefly, images were thresholded to estimate the volume and centroid of individual microparticles. Cubic interpolation was then used to calculate the intensity values along lines originating from the particle centroid and crossing the particle edge. Edge coordinates were then directly localized with super-resolution accuracy by fitting a Gaussian to the discrete derivative of these line profiles. This is significantly faster than using pre-processing of the image stacks with the 3D Sobel operator as used previously^21^. Before calculation of particle properties, such as sphericity, relative elongation and surface curvature, as well as traction forces, edge coordinates were smoothed using the equivalent of a 2D moving average for a spherical surface, operating on the radial component of the edge coordinates. Great circle distances (*d*) between edge coordinates with indices *i* and *j* were calculated along a perfect sphere: *d* = arccos(sin *θ*_*i*_ sin *θ*_*j*_ + cos *θ*_*i*_ cos *θ*_*j*_ cos(*φ*_*i*_ – *φ*_*j*_))*R,* where *R* is the equivalent radius of a sphere to the particle. The radial component of the edge coordinates was then averaged within the given window size (1 μm^2^). A triangulation between edge coordinates was generated, and the particle surface area *S* and volume *V* calculated. Sphericity was calculated as Ψ = (6*π*^1/3^*V*^2/3^*S*^−1^ For surface curvature calculations, first principal curvatures (*k*_1_ and *k*_2_) of the triangulated mesh were determined as described previously^62,63^. The mean curvature was calculated *H* = (*k*_1_ + *k*_2_)/2. Force calculations were performed using the spherical harmonics method within custom Python package ShElastic as described in detail previously^21,64^. Chosen values for the weighing parameter α for residual traction and β for anti-aliasing were both 1.

### Fluorescent mapping on particle surface and determination of fraction engulfed

Mapping of fluorescent proteins, phalloidin and immunostaining to the particle surface was done by determination of the fluorescent intensity along radial lines originating from the particle centroid and passing through each edge coordinate (Fig. 1c). Linear interpolation was used to determine the intensity along each line, and the maximum value within a 1 μm distance of the edge coordinate was projected onto the surface. The calculation of the fraction engulfed, alignment of particles using the centroid of the contact area, and obtaining of a stress-free boundary for force calculations was done as described previously^21^, with the exception that here both the phalloidin stain and the immunostaining of the free particle surface were used to determine the mask (Supplementary Fig. 2a). For LLSM data, where no staining of the free particle surface was present, alignment was done manually.

### Indentation simulations

Indentation simulations of teeth on a spherical target particle were based on the Hertz contact model^65^. Parameters of the model were estimated from experimental data: the undeformed radius of the target particle was set at *R*_target_ = 3.7μm; the teeth are considered as rigid spherical indenters with radius *R*_teeth_ ≈ 0.5 μm; and 10 teeth were simulated for each target, which were equally distributed around the equator of the target sphere (Supplementary Fig. 8a). The force *F* and the contact area radius *a* produced by indentation to absolute depth *d* for each individual indenter were then evaluated:

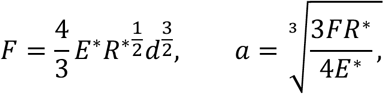

where the effective Young’s modulus *E*\* and effective radius *R*\* are

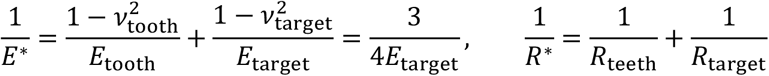

given that the target particle is near incompressible (*ν*_*target*_ = 0.5) and the teeth are rigid compared to the target (*E*_tooth_ ≫ *E*_target_). Considering non-friction contact, the force distribution on the target sphere in the contact area of each tooth can be written as,

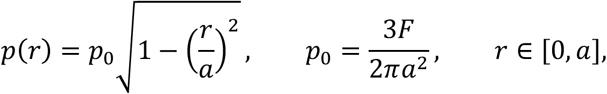

where *r* is the radius to the initial contact point, and *p*_0_ is the maximum pressure on the contact plane. Given the resulting traction force map *T*(*θ*, *φ*) as the boundary condition on the target sphere surface, we solved the elasticity problem, and obtained the displacement map *u*(*θ*, *φ*) using our ShElastic package (Supplementary Fig. 8b)^64^. The (*θ*, *φ*) map on the spherical surface has the size of 61×121, which is defined by Gauss-Legendre quadrature^66^. Simulations were carried out for a range of tooth radii *R*_teeth_ and absolute depth *d* to obtain the effective tooth depth and the average constriction along the equator (Supplementary Fig. 8b), which are directly comparable with experimentally obtained data.

## References

1. Lim, J. J., Grinstein, S. & Roth, Z. Diversity and Versatility of Phagocytosis: Roles in Innate Immunity, Tissue Remodeling, and Homeostasis. Front. Cell. Infect. Microbiol. 7, 1–12 (2017).

2. Boada-Romero, E., Martinez, J., Heckmann, B. L. & Green, D. R. The clearance of dead cells by efferocytosis. Nat. Rev. Mol. Cell Biol. 21, 398–414 (2020).

3. Uribe-Querol, E. & Rosales, C. Phagocytosis: Our Current Understanding of a Universal Biological Process. Front. Immunol. 11, 1–13 (2020).

4. Freeman, S. A. & Grinstein, S. Phagocytosis: receptors, signal integration, and the cytoskeleton. Immunol. Rev. 262, 193–215 (2014).

5. Griffin, F. M. & Silverstein, S. C. Segmental Response of the Macrophage Plasma Membrane to a Phagocytic Stimulus. J. Exp. Med. 139, 323–336 (1974).

6. Swanson, J. A. & Hoppe, A. D. The coordination of signaling during Fc receptor-mediated phagocytosis. J. Leukoc. Biol. 76, 1093–1103 (2004).

7. Jaumouillé, V. & Waterman, C. M. Physical Constraints and Forces Involved in Phagocytosis. Front. Immunol. 11, 1–20 (2020).

8. Vorselen, D., Labitigan, R. L. D. & Theriot, J. A. A mechanical perspective on phagocytic cup formation. Curr. Opin. Cell Biol. 66, 112–122 (2020).

9. Blanchoin, L., Boujemaa-Paterski, R., Sykes, C. & Plastino, J. Actin Dynamics, Architecture, and Mechanics in Cell Motility. Physiol. Rev. 94, 235–263 (2014).

10. Small, J. V., Stradal, T., Vignal, E. & Rottner, K. The lamellipodium: where motility begins. Trends Cell Biol. 12, 112–120 (2002).

11. Davidson, A. J. & Wood, W. Macrophages Use Distinct Actin Regulators to Switch Engulfment Strategies and Ensure Phagocytic Plasticity In Vivo. Cell Rep. (2020). doi:10.1016/j.celrep.2020.107692

12. Barger, S. R. et al. Membrane-cytoskeletal crosstalk mediated by myosin-I regulates adhesion turnover during phagocytosis. Nat Commun 10, 1249 (2019).

13. Ostrowski, P. P., Freeman, S. A., Fairn, G. & Grinstein, S. Dynamic Podosome-Like Structures in Nascent Phagosomes Are Coordinated by Phosphoinositides. Dev. Cell 50, 397–410.e3 (2019).

14. Linder, S. The matrix corroded: podosomes and invadopodia in extracellular matrix degradation. Trends Cell Biol. 17, 107–117 (2007).

15. van den Dries, K., Linder, S., Maridonneau-Parini, I. & Poincloux, R. Probing the mechanical landscape – new insights into podosome architecture and mechanics. J. Cell Sci. 132, jcs236828 (2019).

16. Labernadie, A. et al. Protrusion force microscopy reveals oscillatory force generation and mechanosensing activity of human macrophage podosomes. Nat. Commun. 5, 5343 (2014).

17. van den Dries, K. et al. Modular actin nano-architecture enables podosome protrusion and mechanosensing. Nat. Commun. 10, 1–16 (2019).

18. Beningo, K. A. & Wang, Y. Fc-receptor-mediated phagocytosis is regulated by mechanical properties of the target. J. Cell Sci. 115, 849–56 (2002).

19. Sosale, N. G. et al. Cell rigidity and shape override CD47’s ‘self’-signaling in phagocytosis by hyperactivating myosin-II. Blood 125, 542–552 (2015).

20. Jaumouillé, V., Cartagena-Rivera, A. X. & Waterman, C. M. Coupling of β2 integrins to actin by a mechanosensitive molecular clutch drives complement receptor-mediated phagocytosis. Nat. Cell Biol. 21, 1357–1369 (2019).

21. Vorselen, D. et al. Microparticle traction force microscopy reveals subcellular force exertion patterns in immune cell–target interactions. Nat. Commun. 11, 20 (2020).

22. Swanson, J. A. et al. A contractile activity that closes phagosomes in macrophages. J. Cell Sci. 112 (Pt 3, 307–16 (1999).

23. Araki, N., Hatae, T., Furukawa, A. & Swanson, J. A. Phosphoinositide-3-kinase-independent contractile activities associated with Fcgamma-receptor-mediated phagocytosis and macropinocytosis in macrophages. J Cell Sci 116, 247–257 (2003).

24. Ikeda, Y. et al. Rac1 switching at the right time and location is essential for Fcγ receptor-mediated phagosome formation. J. Cell Sci. 130, 2530–2540 (2017).

25. Kovari, D. T. et al. Frustrated Phagocytic Spreading of J774A-1 Macrophages Ends in Myosin II-Dependent Contraction. Biophys. J. 111, 2698–2710 (2016).

26. Barger, S. R., Gauthier, N. C. & Krendel, M. Squeezing in a Meal: Myosin Functions in Phagocytosis. Trends Cell Biol. 30, 157–167 (2020).

27. Rotty, J. D. et al. Arp2/3 Complex Is Required for Macrophage Integrin Functions but Is Dispensable for FcR Phagocytosis and In Vivo Motility. Dev. Cell 42, 498–513.e6 (2017).

28. Olazabal, I. M. et al. Rho-kinase and myosin-II control phagocytic cup formation during CR, but not FcgammaR, phagocytosis. Curr Biol 12, 1413–1418 (2002).

29. Heinrich, V. Controlled One-on-One Encounters between Immune Cells and Microbes Reveal Mechanisms of Phagocytosis. Biophys. J. 109, 469–476 (2015).

30. Herant, M. Mechanics of neutrophil phagocytosis: behavior of the cortical tension. J. Cell Sci. 118, 1789–1797 (2005).

31. Rougerie, P. & Cox, D. Spatio-temporal mapping of mechanical force generated by macrophages during FcγR-dependent phagocytosis reveals adaptation to target stiffness. bioRxiv (2020). doi:10.1101/2020.04.14.041335

32. Mohagheghian, E. et al. Quantifying compressive forces between living cell layers and within tissues using elastic round microgels. Nat. Commun. 9, 1878 (2018).

33. Träber, N. et al. Polyacrylamide Bead Sensors for in vivo Quantification of Cell-Scale Stress in Zebrafish Development. Sci. Rep. 9, 17031 (2019).

34. Horsthemke, M. et al. Multiple roles of filopodial dynamics in particle capture and phagocytosis and phenotypes of Cdc42 and Myo10 deletion. J. Biol. Chem. 292, 7258–7273 (2017).

35. Freeman, S. A. et al. Lipid-gated monovalent ion fluxes regulate endocytic traffic and support immune surveillance. Science (80-.). (2020). doi:10.1126/science.aaw9544

36. Liebl, D. & Griffiths, G. Transient assembly of F-actin by phagosomes delays phagosome fusion with lysosomes in cargo-overloaded macrophages. J. Cell Sci. (2009). doi:10.1242/jcs.048355

37. Legant, W. R. et al. Multidimensional traction force microscopy reveals out-of-plane rotational moments about focal adhesions. Proc. Natl. Acad. Sci. 110, 881–886 (2013).

38. Case, L. B. & Waterman, C. M. Integration of actin dynamics and cell adhesion by a three-dimensional, mechanosensitive molecular clutch. Nat. Cell Biol. 17, 955–963 (2015).

39. van Zon, J. S., Tzircotis, G., Caron, E. & Howard, M. A mechanical bottleneck explains the variation in cup growth during FcgammaR phagocytosis. Mol. Syst. Biol. 5, 298 (2009).

40. Wilson, C. A. et al. Myosin II contributes to cell-scale actin network treadmilling through network disassembly. Nature 465, 373–377 (2010).

41. Ostrowski, P. P., Freeman, S. A., Fairn, G. & Grinstein, S. Dynamic Podosome-Like Structures in Nascent Phagosomes Are Coordinated by Phosphoinositides. Dev. Cell 50, 397–410.e3 (2019).

42. Proag, A. et al. Working together: Spatial synchrony in the force and actin dynamics of podosome first neighbors. ACS Nano 9, 3800–3813 (2015).

43. Meddens, M. B. M. et al. Actomyosin-dependent dynamic spatial patterns of cytoskeletal components drive mesoscale podosome organization. Nat. Commun. 7, 1–17 (2016).

44. Greenberg, S., Burridge, K. & Silverstein, S. C. Colocalization of F-actin and talin during Fc receptor-mediated phagocytosis in mouse macrophages. J Exp Med 172, 1853–1856 (1990).

45. Herant, M., Heinrich, V. & Dembo, M. Mechanics of neutrophil phagocytosis: experiments and quantitative models. J. Cell Sci. 119, 1903–13 (2006).

46. Tollis, S., Dart, A. E., Tzircotis, G. & Endres, R. G. The zipper mechanism in phagocytosis: energetic requirements and variability in phagocytic cup shape. BMC Syst. Biol. 4, 149 (2010).

47. Matlung, H. L. et al. Neutrophils Kill Antibody-Opsonized Cancer Cells by Trogoptosis. Cell Rep. 23, 3946–3959.e6 (2018).

48. Velmurugan, R., Challa, D. K., Ram, S., Ober, R. J. & Ward, E. S. Macrophage-mediated trogocytosis leads to death of antibody-opsonized tumor cells. Mol. Cancer Ther. 15, 1879–1889 (2016).

49. Morrissey, M. A. et al. Chimeric antigen receptors that trigger phagocytosis. Elife 7, e36688 (2018).

50. Walbaum, S. et al. Complement receptor 3 mediates both sinking phagocytosis and phagocytic cup formation via distinct mechanisms. J. Biol. Chem. 296, 100256 (2021).

51. Hoijman, E. et al. Cooperative epithelial phagocytosis enables error correction in the early embryo. Nature 590, 618–623 (2021).

52. Bieling, P. et al. Force Feedback Controls Motor Activity and Mechanical Properties of Self-Assembling Branched Actin Networks. Cell 164, 115–127 (2016).

53. Marie-Anaïs, F., Mazzolini, J., Bourdoncle, P. & Niedergang, F. ‘Phagosome Closure Assay’ to Visualize Phagosome Formation in Three Dimensions Using Total Internal Reflection Fluorescent Microscopy (TIRFM). J. Vis. Exp. 2016, 1–8 (2016).

54. Nishi, H. et al. Neutrophil FcγRIIA promotes IgG-mediated glomerular neutrophil capture via Abl/Src kinases. J. Clin. Invest. 127, 3810–3826 (2017).

55. Zhu, C., Chen, W., Lou, J., Rittase, W. & Li, K. Mechanosensing through immunoreceptors. Nat. Immunol. 20, 1269–1278 (2019).

56. Schuerle, S. et al. Robotically controlled microprey to resolve initial attack modes preceding phagocytosis. Sci. Robot. 2, eaah6094 (2017).

57. Champion, J. A. & Mitragotri, S. Role of target geometry in phagocytosis. Proc. Natl. Acad. Sci. U. S. A. 103, 4930–4 (2006).

58. Bakalar, M. H. et al. Size-Dependent Segregation Controls Macrophage Phagocytosis of Antibody-Opsonized Targets. Cell 174, 131–142.e13 (2018).

59. Colucci-Guyon, E., Tinevez, J.-Y., Renshaw, S. A. & Herbomel, P. Strategies of professional phagocytes in vivo: unlike macrophages, neutrophils engulf only surface-associated microbes. J. Cell Sci. 124, 3053–3059 (2011).

60. Gosavi, R. A. et al. Optimization of Ex Vivo Murine Bone Marrow Derived Immature Dendritic Cells: A Comparative Analysis of Flask Culture Method and Mouse CD11c Positive Selection Kit Method. Bone Marrow Res. (2018). doi:10.1155/2018/3495086

61. Chen, B. C. et al. Lattice light-sheet microscopy: Imaging molecules to embryos at high spatiotemporal resolution. Science (80-.). (2014). doi:10.1126/science.1257998

62. Rusinkiewicz, S. Estimating curvatures and their derivatives on triangle meshes. in Proceedings. 2nd International Symposium on 3D Data Processing, Visualization and Transmission, 2004. 3DPVT 2004. 486–493 (IEEE, 2004). doi:10.1109/TDPVT.2004.1335277

63. Ben Shabat, Y. & Fischer, A. Design of Porous Micro-Structures Using Curvature Analysis for Additive-Manufacturing. Procedia CIRP 36, 279–284 (2015).

64. Wang, Y., Zhang, X. & Cai, W. Spherical harmonics method for computing the image stress due to a spherical void. J. Mech. Phys. Solids 126, 151–167 (2019).

65. Johnson, K. L. Contact Mechanics. (Cambridge University Press, 1985).

66. Wieczorek, M. A. & Meschede, M. SHTools: Tools for Working with Spherical Harmonics. Geochemistry, Geophys. Geosystems 19, 2574–2592 (2018).

