## Supplementary Information for "Phagocytic “teeth” and myosin-II “jaw” power target constriction during phagocytosis"

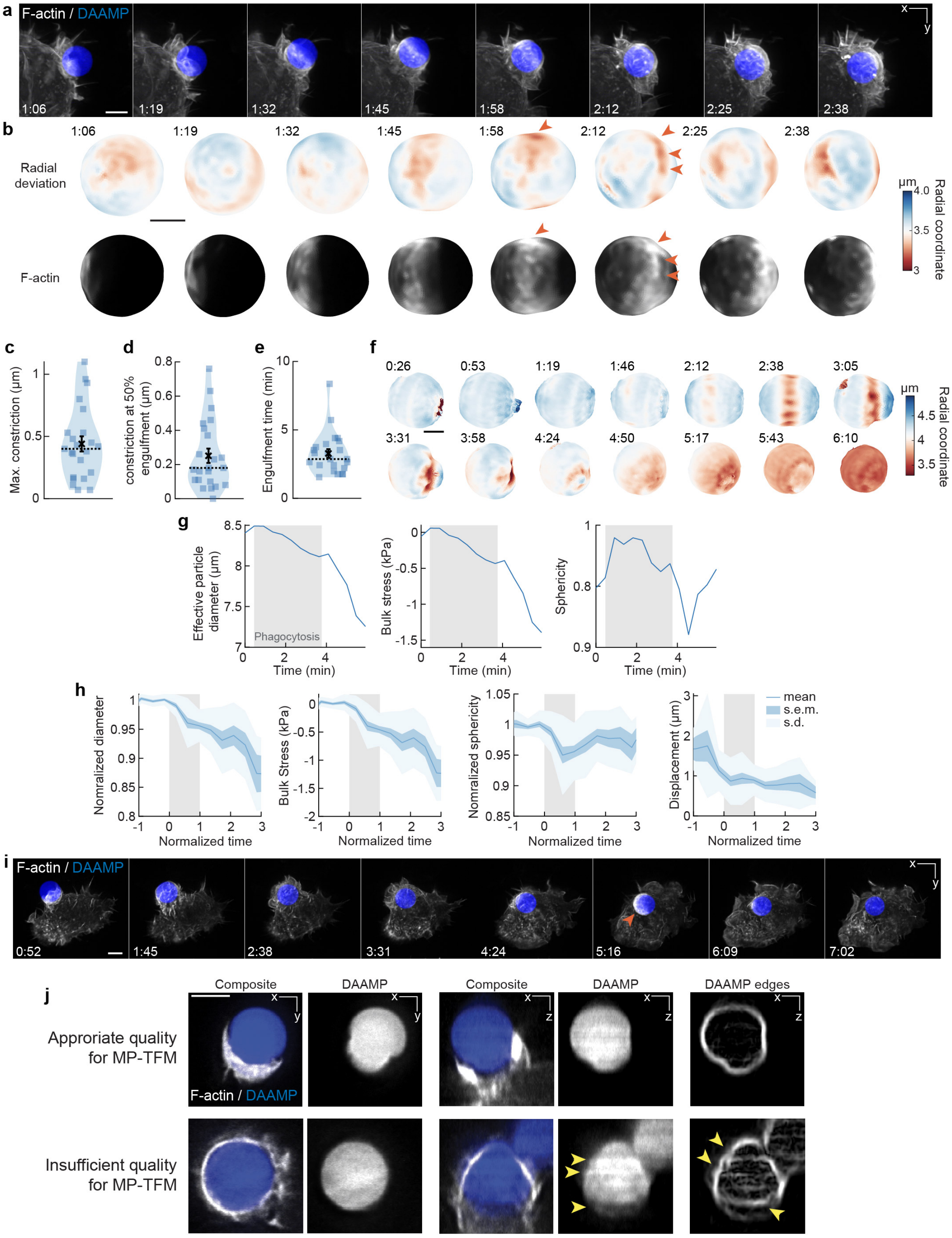


**Supplementary Figure 1**. **Phagocytosis involves similar mechanical sequence for stiffer targets as soft ones, and bulk compressive stresses arise during phagosome formation and maturation. a**, Time lapse montage (min:s) of RAW macrophage transfected with mEmerald-

Lifeact internalizing a DAAMP (9 μm, Young's modulus 6.5 kPa) functionalized with BSA and anti-BSA IgG and AF647-Cadaverine and imaged using lattice light-sheet microscopy (LLSM). Maximum intensity projections in x/y are shown. Scale bar, 5 μm **b,** Side view of reconstructed DAAMP internalized in **a** showing target deformations and F-actin localization on particle surface. Arrows point at loci of F-actin accumulation and protrusion into the target, similar to those observed for 1.4 kPa particles (Fig. 1b). Scale bar, 3 μm. **c,d,e**, Violin plot shows individual phagocytic events (colored markers, n = 23), mean (black cross) and median (dashed line). **c,** Maximum target constriction for live cell uptake movies **d,** Average target constriction at 50% engulfment. Average is 0.25 ± 0.04 μm (s.e.m., n = 23 cups), which is ~40% higher than observed with fixed cell data at similar stage at 0.18 ± 0.02 μm (s.e.m., n = 19). This suggests that the fixed cell force measurements are a slight underestimate of the real phagocytic forces. **e,** Engulfment time for individual phagocytic events. **f,** Volume of DAAMPs decreases with phagocytic internalization. Since hydrogel microparticles are not completely incompressible, their volume can decrease under exertion of bulk forces[1]. Same event is shown as in Fig. 1a. Color scale denotes radial distance to the centroid. Time stamps are provided in min:s, and internalization is complete at the 3:58 time point. Scale bar, 3 μm. **g,** Quantification of effective particle diameter, bulk stress and sphericity over time for DAAMP in **f**. Bulk compressive forces can be estimated from the previous DAAMP bulk modulus measurements (~3.8 kPa)[1]. Grey area indicates time interval of phagocytosis. Compressive stresses arise during phagocytosis and are increased after completion of internalization. **h,** Quantification of bulk stresses, target sphericity and displacement of 23 live cell phagocytic events. Compressive stresses are exerted during phagocytosis (~0.5 kPa) and increase after completion (~1.3 kPa). Particle sphericity dips during phagocytosis, but the particles return to a more spherical shape after internalization completion. Grey area indicates the duration of phagocytosis, where normalized time t = 0 indicates the start of the phagocytic event, and t = 1 internalization completion for individual events. Particle diameter and sphericity (because it could be strongly affected by imaging artifacts, see **j**) were normalized to 1, using the measurements before the start of phagocytosis. **i**, Brief F-actin accumulation (orange arrow) observed on DAAMP phagosome following internalization. Time lapse montage (min:s) of RAW macrophage transfected with mEmerald-Lifeact internalizing a DAAMP (9 μm, 6.5 kPa) functionalized with BSA and anti-BSA IgG and AF647-Cadaverine and imaged using lattice light-sheet microscopy (LLSM). Scale bar, 5 μm **j,** Lattice light sheet microscopy image artifacts hinder particle reconstruction. Top row, a xy- and xz-slice through a (rare) particle that is barely affected by artifacts. Bottom row, typical images strongly affected by artifacts. Artifacts are not obvious in the xy-plane, but a strong “striping” artifact, with strong fluctuations in fluorescent intensity (yellow arrows) are visible along the optical axis. Shape reconstruction in MP-TFM is critically dependent on particle edge localization detection, which shows clear irregularities (yellow arrows) because of the striping artifact. Scale bar: 5 μm.

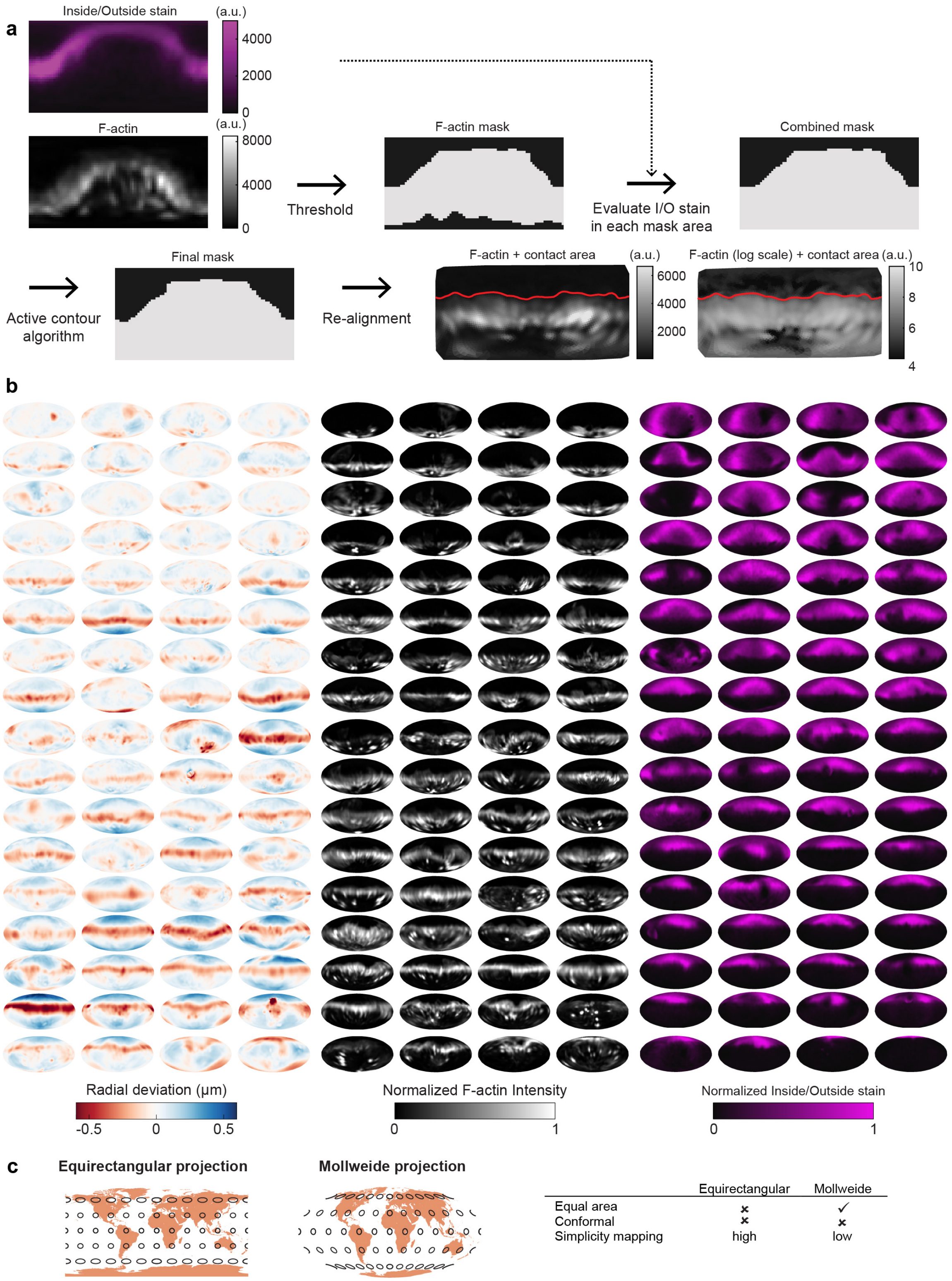


**Supplementary Figure 2. Automated image analysis reveals deformation, F-actin localization, and Inside/Outside stain for 68 phagocytic events. a,** Determination of contact area between cell and target was calculated using both inside/outside (I/O) signal (immunostain of exposed particle surface) and the F-actin signal for confocal microscopy data. First, an initial mask was determined based on thresholding the logarithm of the F-actin signal. Then, for each area with low F-actin intensity the I/O stain is compared to the mean I/O stain, and each area is marked as inside (when the I/O stain is below average) or outside (when the I/O stain is above average). A region-based active contour algorithm was used to optimize the mask[1,2]. Finally, particles were aligned, and their fraction engulfed (masked area/total area) calculated. **b,** Visualization of deformations, F-actin intensity and I/O stain for all 68 control particles (being phagocytosed by DMSO-treated cells) used in this manuscript shown using Mollweide projections (see **c**). **c,** Comparison of Equirectangular projection and Mollweide projection for mapping spherical surface for 2D visualization. All 2D projections of a sphere are necessarily distorted. The Mollweide projection is an equal area projection, and therefore does not lay emphasis on the polar regions. Angles are, however, not preserved in this mapping.

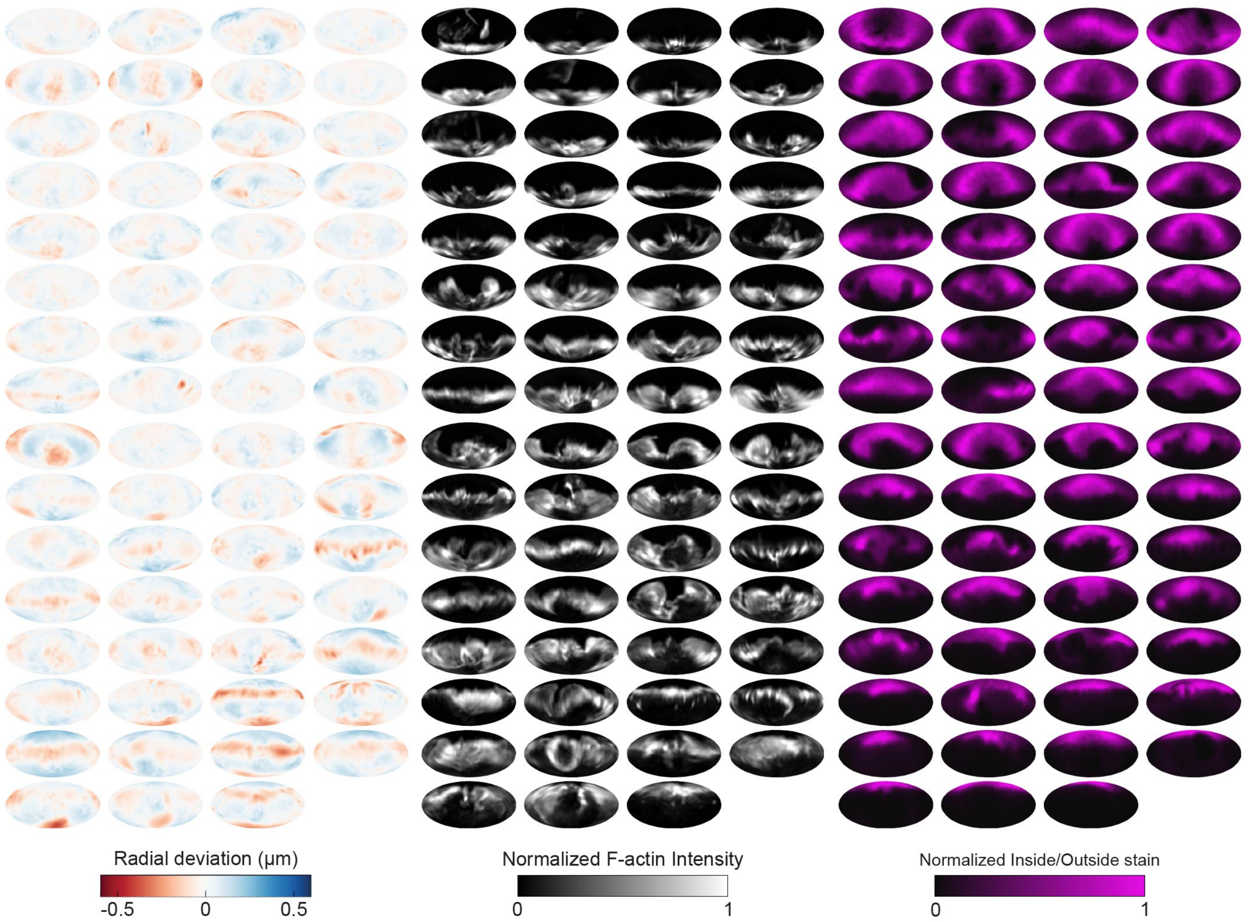


**Supplementary Figure 3**. **Automated image analysis reveals deformation, F-actin localization, and Inside/Outside stain for 63 phagocytic events in CK666-treated cells.** Visualization of deformations, F-actin intensity and I/O stain for all 63 particles being phagocytosed by CK666-treated cells used in this manuscript shown using Mollweide projections.

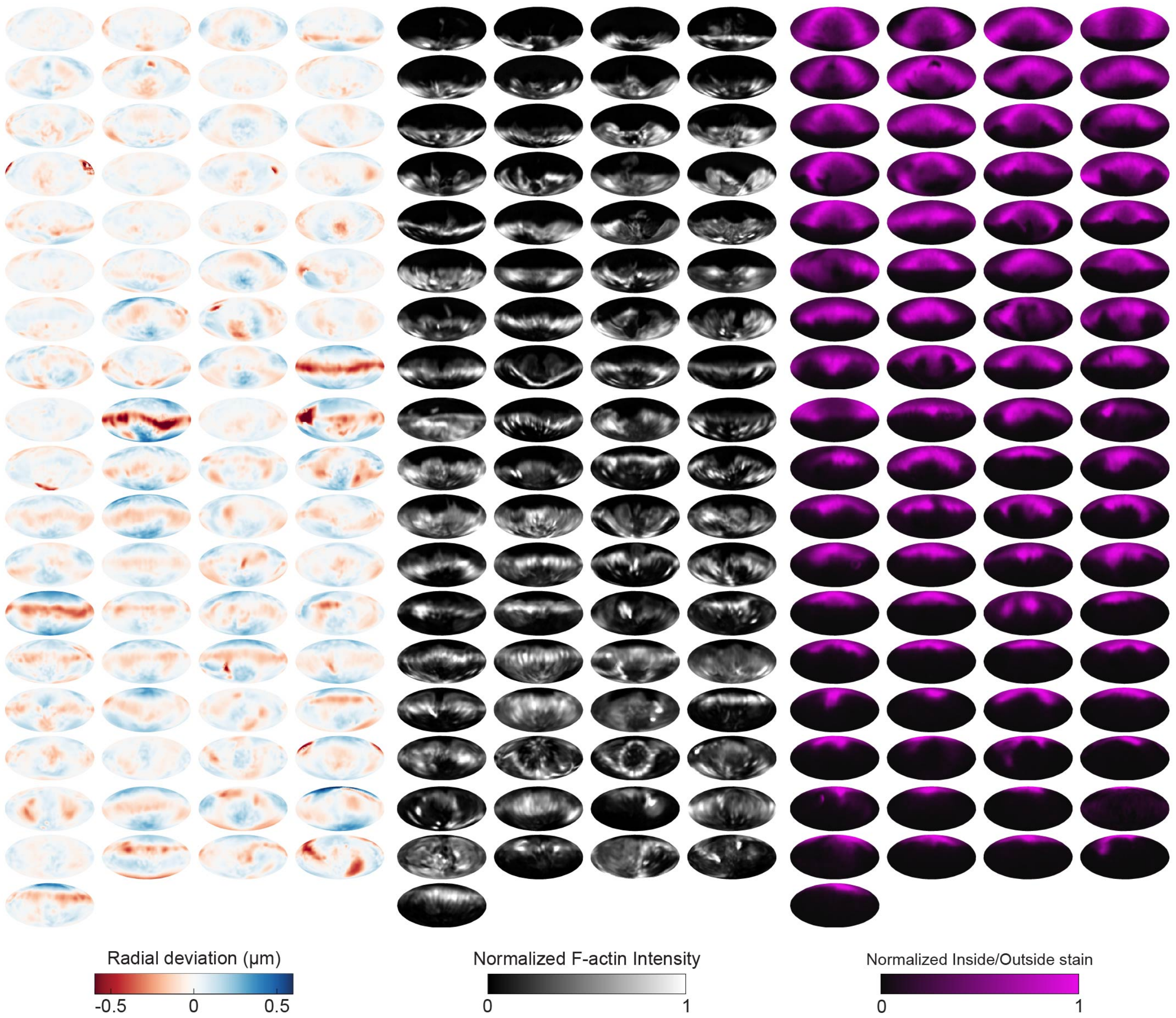


**Supplementary Figure 4**. **Automated image analysis reveals deformation, F-actin localization, and Inside/Outside stain for 75 phagocytic events in blebbistatin-treated cells.** Visualization of deformations, F-actin intensity and I/O stain for all 75 particles being phagocytosed by blebbistatin-treated cells used in this manuscript shown using Mollweide projections.

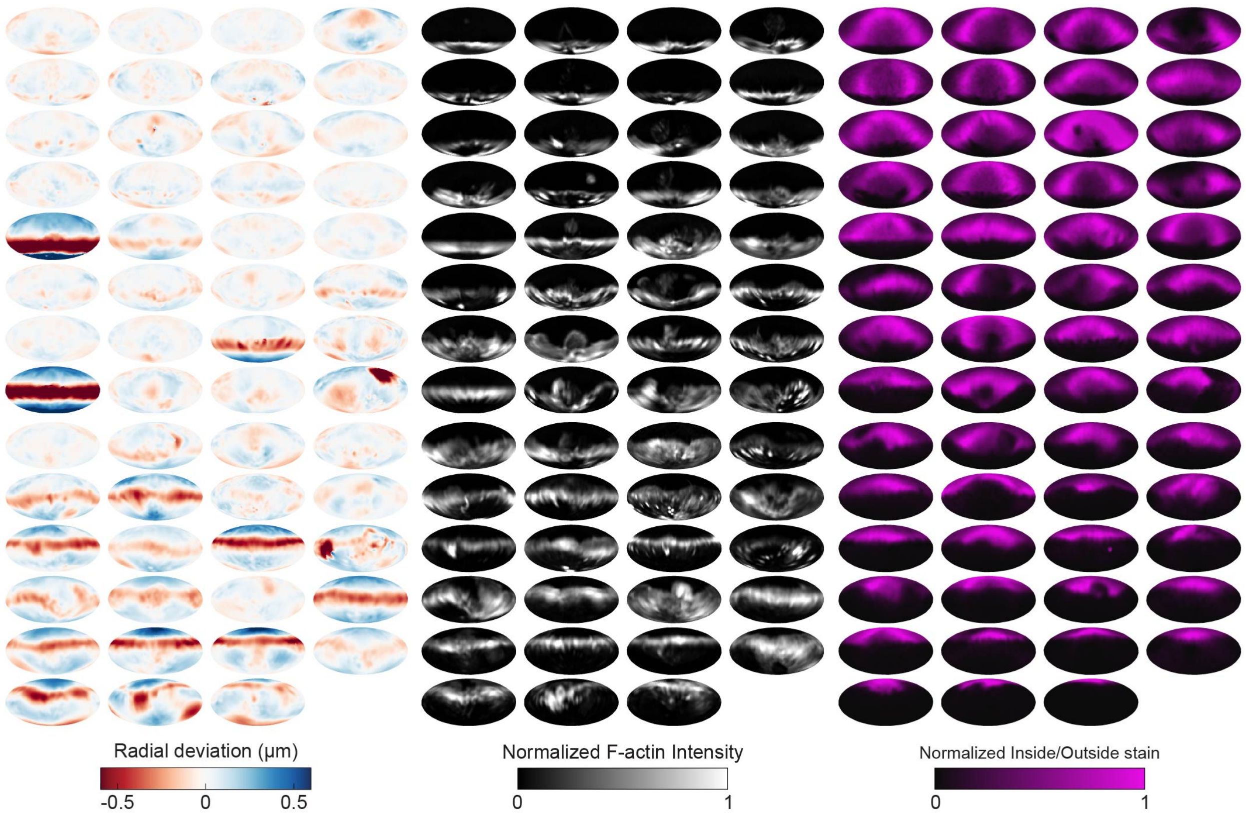


**Supplementary Figure 5**. **Automated image analysis reveals deformation, F-actin localization, and Inside/Outside stain for 55 phagocytic events in SMIFH2-treated cells.** Visualization of deformations, F-actin intensity and I/O stain for all 55 particles being phagocytosed by SMIFH2-treated cells used in this manuscript shown using Mollweide projections.

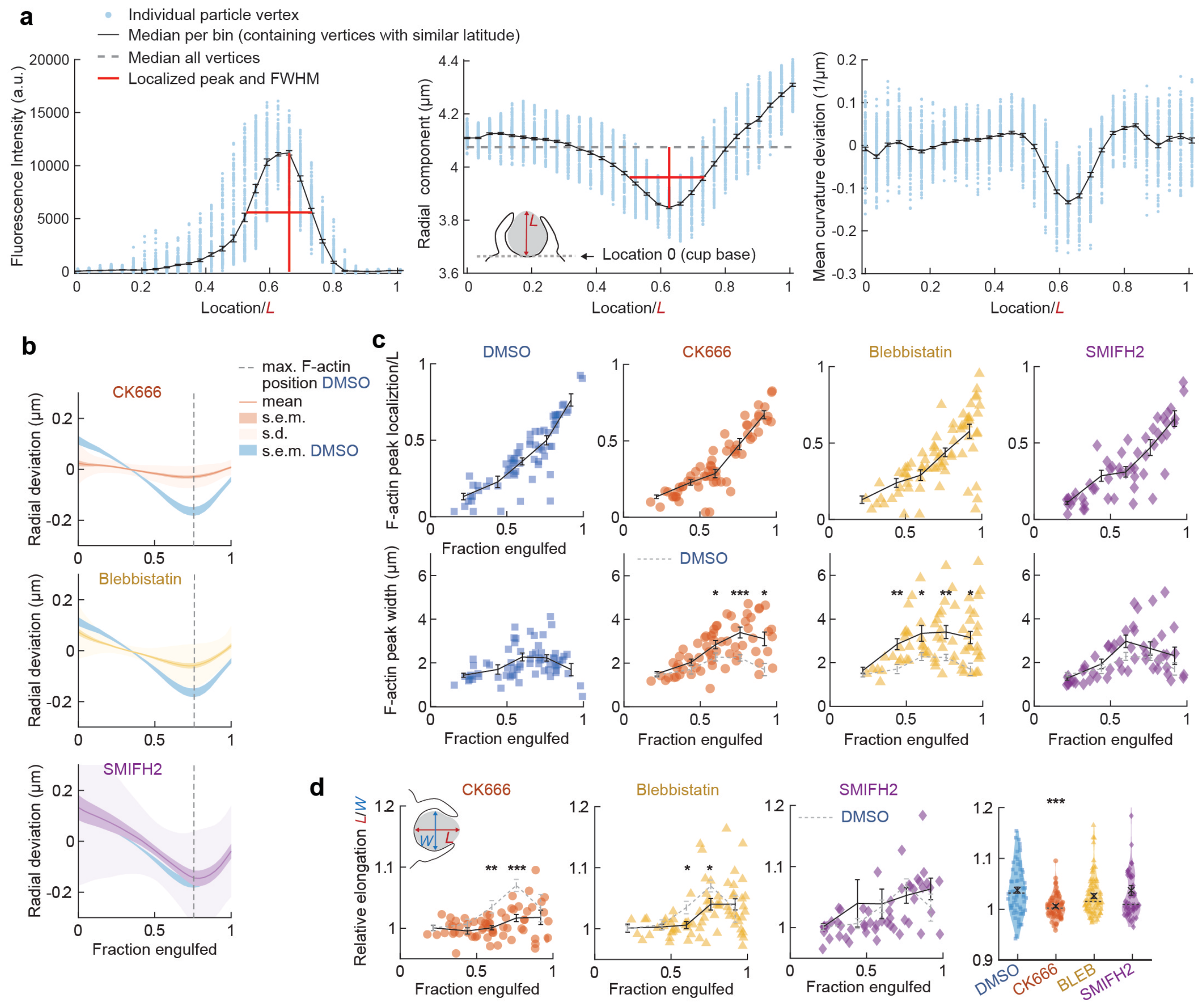


**Supplementary Figure 6**. **Actin organization and target deformation along the phagocytic axis are affected by perturbation of actomyosin activity. a,** Determination of F-actin and contractile ring localization, peak magnitude and peak width. Particles were aligned with the centroid of the cell-target contact area (base of the cup) at the south pole (see inset middle panel). All particle edge coordinates (~4250) were then divided into bins per latitude of which the median was calculated. Finally, peak height, location and full width at half maximum (FWHM) were determined. Data shown corresponds to the particle in the 3$^{rd}$ column, 2$^{nd}$ to last row in Supplementary Fig. 2b. **b,** Average target deformation profiles along the phagocytic axis until the cup rim. Signals were first processed on a per-particle basis by averaging over the surface along the phagocytic targets in 30 bins. **c,** Determination of F-actin statistics along the phagocytic axis for all phagocytic events (n = 68, 63, 73 & 55 respectively). Colored markers indicate individual measurements, black lines indicate averages within 5 bins. All statistical tests were two-side Wilcoxon rank sum test comparing with the DMSO control over the same bin with significance levels: $p < 0.05$*; $p < 0.01$**; $p < 0.001$***, unless otherwise indicated. **d,** Target elongation changes with phagocytic progression upon drug treatment. Marker and line styles as in **c**. Inset in leftmost panel shows schematically how relative elongation was determined. Right column, violin plots of all events, showing individual phagocytic events (colored markers) mean (black cross), median (dashed line).

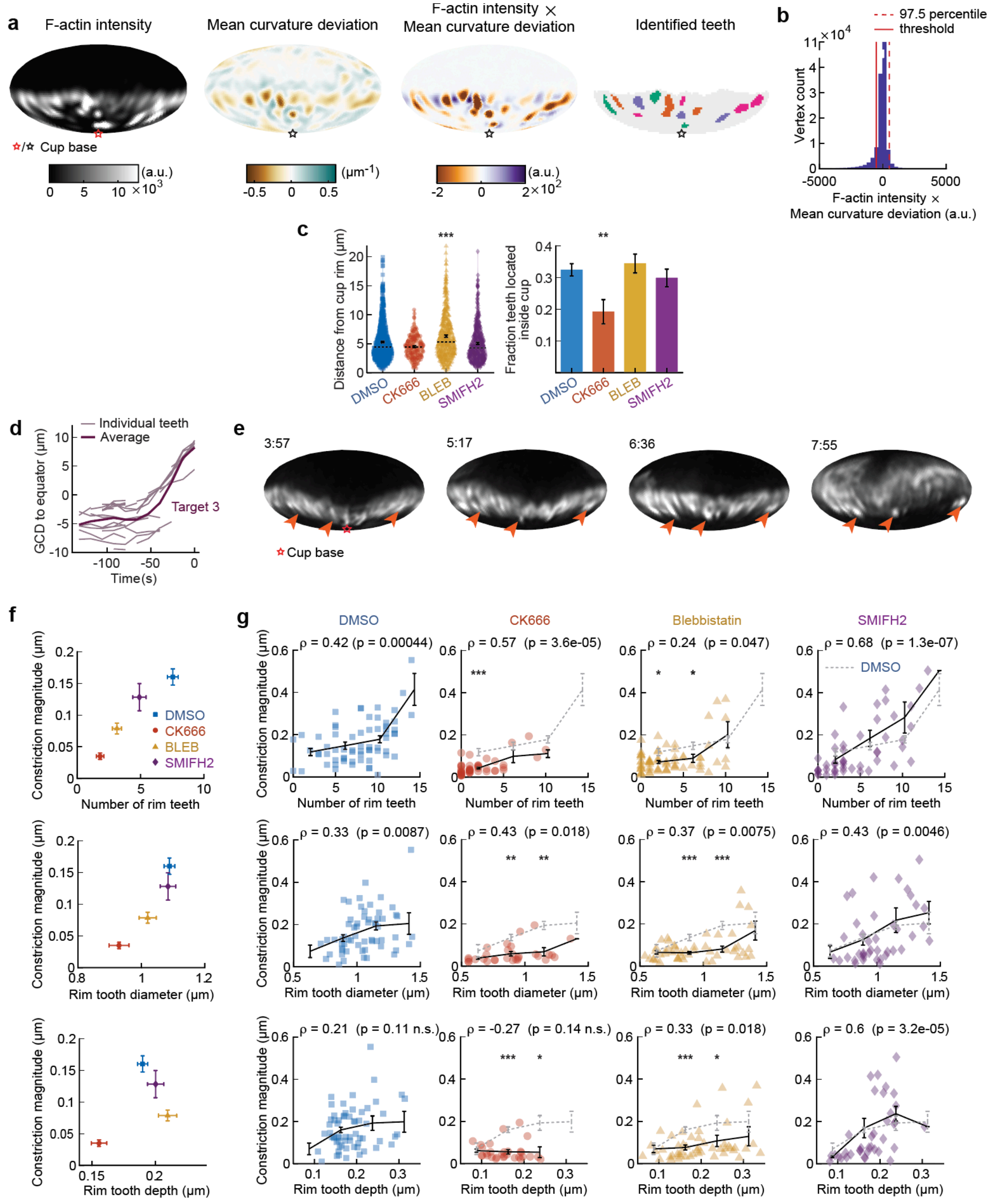


**Supplementary Figure 7. Automated teeth identification reveals how teeth physical properties correlate with overall target constriction. a,** Example of teeth identification method. Teeth were defined as areas of high-actin intensity and inward protrusion. For the mean curvature, the average value of mean curvature for edge coordinates (vertices) at similar latitude (with the base of the cup at the south pole) were subtracted to negate any influence of the typical ring of target constriction at the rim of the cup. Then the values of corrected mean curvature deviation and F-actin intensity were multiplied for each vertex. Vertices with values above the threshold were identified as teeth, and a watershed algorithm was used to separate clusters of teeth. Base of the phagocytic cup is at the south pole. **b,** Determination of the threshold for tooth identification base on the product of F-actin intensity and mean curvature deviation. Generally, there is a negative correlation between F-actin intensity and mean curvature (high F-actin concentration is found at regions of indentation). The threshold was set as the negative of the 97.5 percentile value, which represents an estimate of the mean - 2 standard deviations in the case that F-actin would not have correlated with inward deformation. **c,** Actomyosin activity affects teeth localization. Markers represent individual teeth (n = 716, 138, 377 & 363 from left to right) for all recorded phagocytic events (n = 68, 63, 73 & 55, respectively). Violin plots show individual teeth (colored markers), means (black cross), median (dashed line). Right; fraction of teeth localized in the rim as opposed to deeper in the cup. The threshold for differentiating teeth in the rim from teeth throughout the cup was based on the distribution of teeth distances from the rim in the DMSO condition. Specifically, it was set as the mode of the distribution (see left panel) summed with 2 x the 2.5 percentile value of the distribution, which represents an estimate of the mean tooth distance from the cup edge - 2 standard deviations of the teeth specifically localized in the rim. Fisher's exact test was used to compare fractions ($p = 0.0076$**). Error bars indicate st.d. estimated by treating phagocytosis as a Bernoulli process. **d,** Great circle distance (GCD) of manually tracked teeth to the equator for a third phagocytic event. Data of the other phagocytic events is presented are presented in figure 4i. **e,** Time lapse montage (min:s) showing example of stationary F-actin teeth during phagocytosis. F-actin signal reconstructed over the target surface from LLSM data. Particle surface is shown using Mollweide projection (Supplementary Fig. 2c). Four timepoints of a single phagocytic event are shown. Arrows point at locations of F-actin puncta that can be tracked from an early stage and are still present at the same location in late-stage phagocytosis, when they are far behind the rim of the phagocytic cup. **f,** Number of rim teeth and tooth diameter variation between drug treatment groups correlates with constriction magnitude. **g,** Teeth statistics correlate with constriction magnitude within drug treatment groups. Markers indicate individual measurements; black lines indicate averages within 5 bins. Spearman's rank correlation coefficient (r) is given above each graph. All statistical tests were two-side Wilcoxon rank sum test comparing with the DMSO control over the same bin with significance levels: $p < 0.05$*; $p < 0.01$**; $p < 0.001$***. All error bars indicate s.e.m.

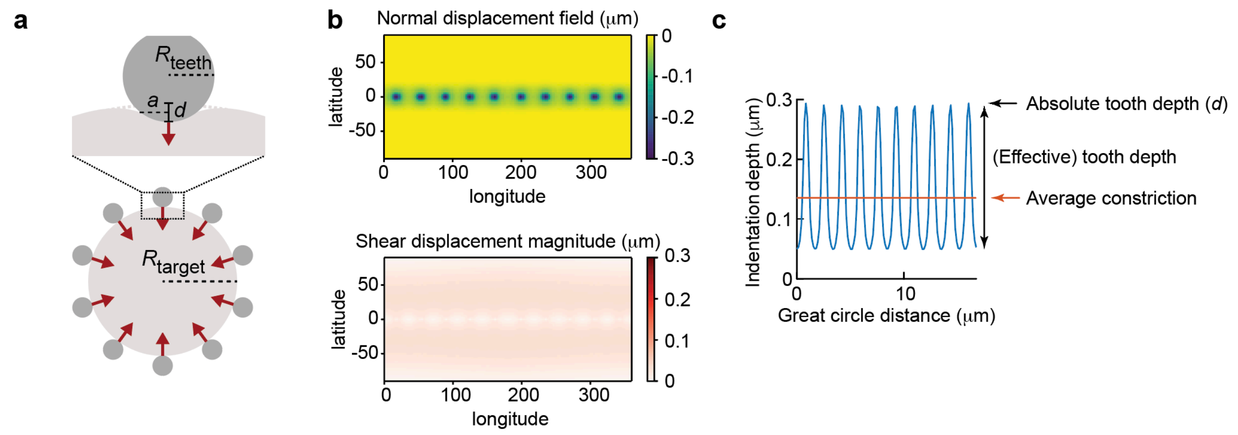


**Supplementary Figure 8. Indentation simulations of phagocytic teeth allow comparison of teeth physical properties and total target constriction. a,** Parametrization of the model. 10 rigid teeth indent a target around the equator of the particle, with $R_{teeth}$ the tooth radius, $R_{target}$ the target radius (3.7 μm), $d$ the absolute indentation depth, and $a$ the contact radius. Red arrows indicate direction of simulated forces. **b**, Displacement result for $d$ = 0.3 μm, $R_{tooth}$ = 0.5 μm. **c,** Derivation of quantities that are obtained from experimental data from the simulation results. The indentation depth variation is shown along the equator of the particle in blue, for the same parameters as **b**. Averaging this profile allows us to find the average constriction depth. The effective tooth depth is then found by subtracting the minimum indentation from the maximum indentation.

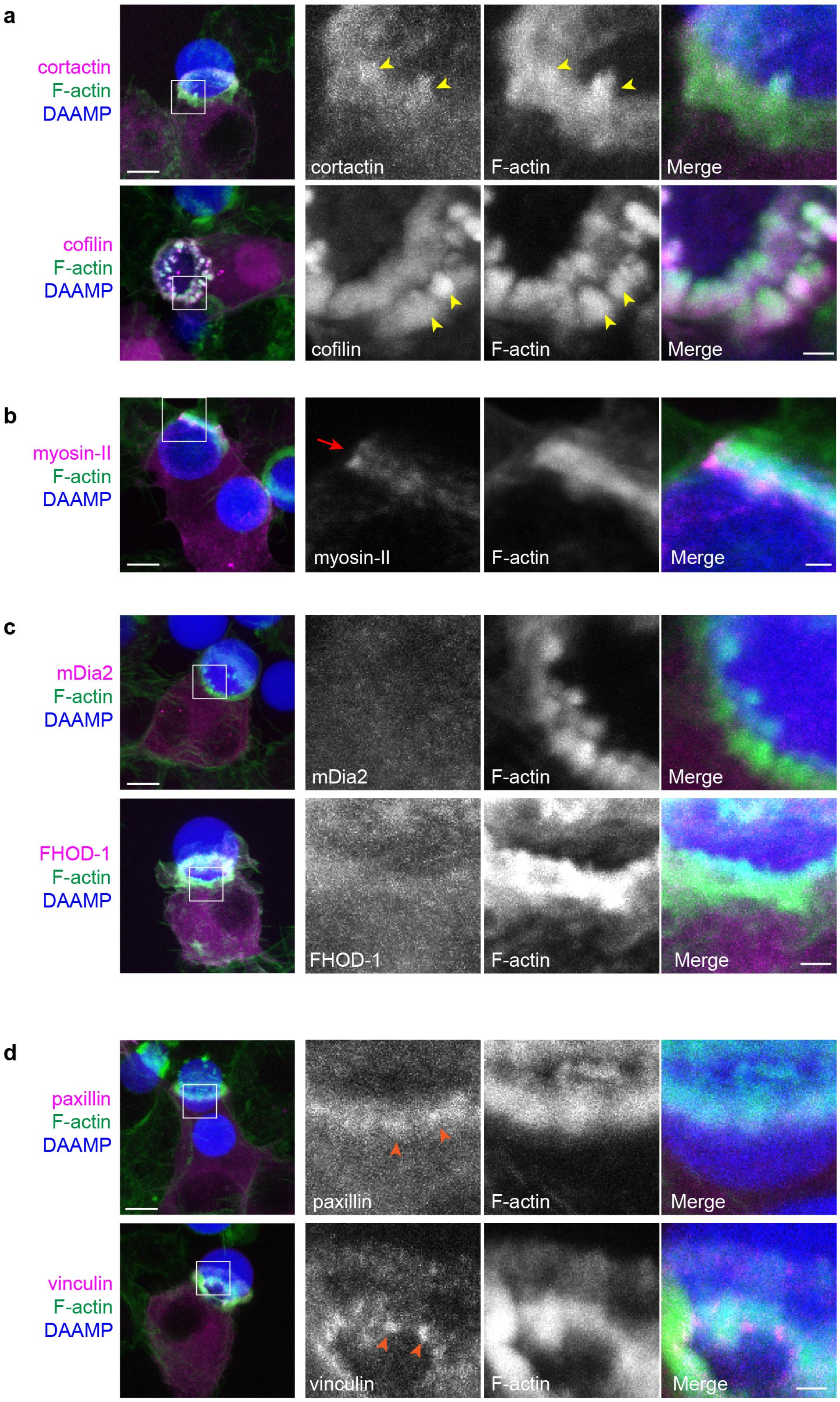

a
cortactin
F-actin
DAAMP
cortactin
F-actin
Merge
cofilin
F-actin
DAAMP
cofilin
F-actin
Merge
b
myosin-II
F-actin
DAAMP
myosin-II
F-actin
Merge
c
mDia2
F-actin
DAAMP
mDia2
F-actin
Merge
FHOD-1
F-actin
DAAMP
FHOD-1
F-actin
Merge
d
paxillin
F-actin
DAAMP
paxillin
F-actin
Merge
vinculin
F-actin
DAAMP
vinculin
F-actin
Merge

**Supplementary Figure 9. Multiple actin-binding proteins localize to phagocytic teeth.** RAW macrophages were transfected with fluorescently tagged actin binding proteins and challenged to ingest DAAM-particles (11 μm, 1.4 kPa) functionalized with AF647-Cadaverine, BSA and anti-BSA IgG. **a,** Actin-binding proteins, cortactin and cofilin localize to phagocytic teeth (yellow arrows) during DAAM-particle internalization **b,** Myosin-II forming concentric rings (marked by red arrow) during late-stage phagocytosis. **c,** Formins, mDia2 and FHOD1, do not distinctly localize to the actin teeth of the phagocytic cup during DAAM-particle internalization. **d,** Adaptor proteins, vinculin and paxillin, localize behind phagocytic teeth in a punctate pattern (orange arrows). DAAMPs are 9 μm, 1 kPa. Images are maximum intensity projections of confocal Z-stacks. Scale bar, 5 μm. Zoom scale bar, 1 μm.

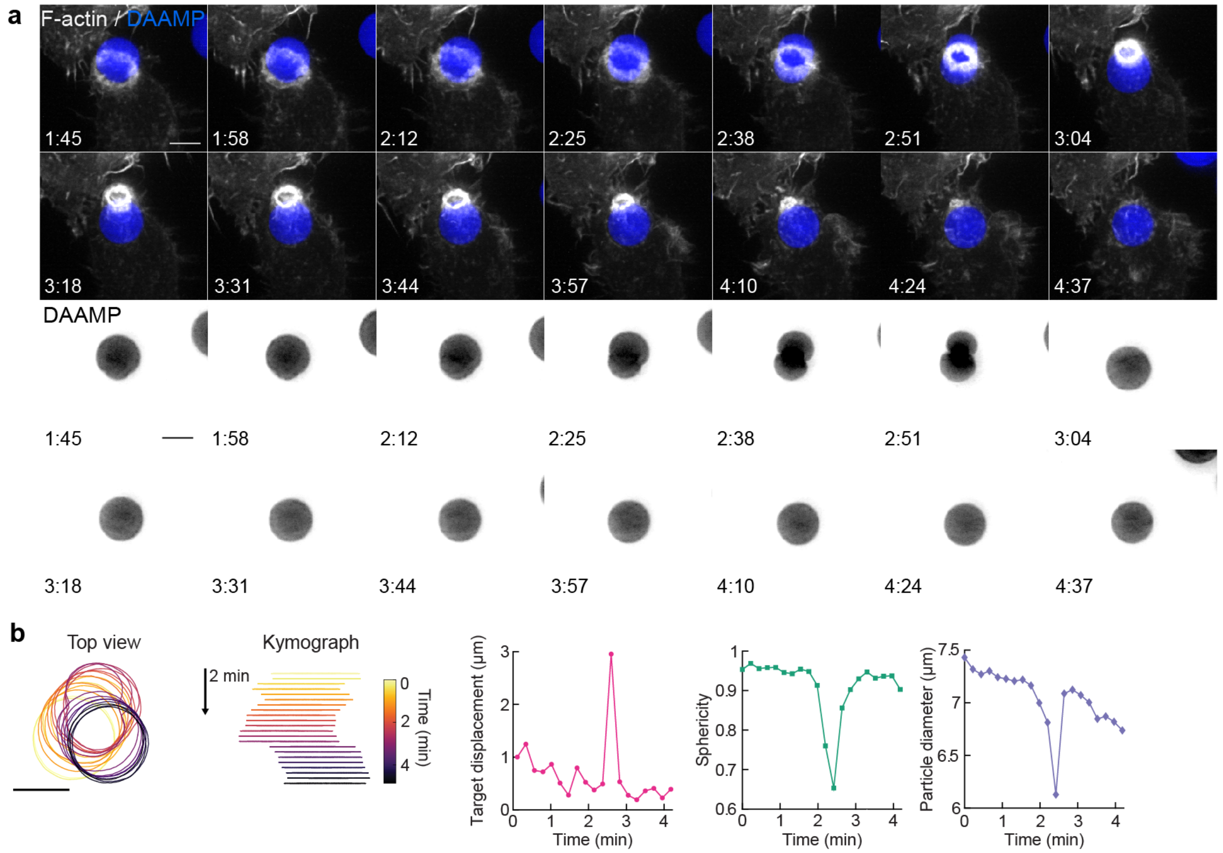


**Supplementary Figure 10. Target squeezing can result in the target popping into the phagocytic cup. a,** Top, maximum intensity projections (MIP) of LLSM time lapse (min:s) showing successful internalization attempt of RAW macrophage with IgG-functionalized 1.4 kPa DAAM-particle, showing strong deformation and a sudden internalization step. Bottom, DAAM-particle only channel (inverted grayscale) **b,** Left, particle position and outline with color-coded kymograph of particle position. Right, particle displacement, sphericity and apparent diameter over time of the same event shows the sudden nature of the internalization. Scale bars, 5 µm.

**Supplementary Movie 1. RAW macrophage ingesting DAAMP imaged by lattice light-sheet microscopy.** RAW macrophages transfected with mEmerald-Lifeact (gray) were fed DAAM-particles (9 μm,1.4 kPa) (blue) functionalized with AF647-Cadaverine, BSA and anti-BSA IgG. Maximum intensity projections in xy (left) and xz (right). Lower left time stamp: min:s. Scale bar, 5 μm.

**Supplementary Movie 2. 3D microparticle shape reconstruction shows phagocytic force-induced deformations in real time.** Base, side, and front views of reconstructed DAAMP internalized in Fig. 1a & Supplementary Movie 1 showing target deformations (above) and F-actin localization on particle surface (below). Color-scales for radial deviation and F-actin intensity shown on right. Upper right time stamp: min:s. Scale bar, 3 μm.

**Supplementary Movie 3. RAW macrophage ingesting stiffer DAAMP shows similar phagocytic force induced deformation patterns by lattice light sheet microscopy.** Maximum intensity projection of RAW macrophage transfected with mEmerald-Lifeact (gray) ingesting DAAM-particles (9 μm, 6.5 kPa) (blue) functionalized with AF647-Cadaverine, BSA and anti-BSA IgG. Lower left time stamp: min:s. Scale bar, 5 μm.

**Supplementary Movie 4. DAAMP phagosome briefly accumulates F-actin, imaged by lattice light sheet microscopy.** Maximum intensity projection of RAW macrophage transfected with mEmerald-Lifeact (gray) ingesting DAAM-particles (9 μm, 6.5 kPa) (blue) functionalized with AF647-Cadaverine, BSA and anti-BSA IgG. Lower left time stamp: min:s. Scale bar, 5 μm.

**Supplementary Movie 5. Failed DAAMP phagocytosis imaged by lattice light-sheet microscopy.** Maximum intensity projections of RAW macrophages transfected with mEmerald-Lifeact (gray) challenged with DAAM-particles (9 μm,1.4 kPa) (blue). Merged images (left) with single DAAMP channel in gray (right) to highlight target deformations. Time stamp: min:s. Scale bar, 5 μm.

**Supplementary Movie 6. Primary BMDM partakes in DAAMP meal sharing imaged by lattice light-sheet microscopy.** Murine bone-marrow derived macrophages (BMDM) transfected with mEmerald-Lifeact (gray) were fed DAAM-particles (11 μm,1.4 kPa) (blue) functionalized with TRITC-Cadaverine, BSA and anti-BSA IgG. Merged maximum intensity projections (left) with single DAAMP channel in gray (right) to highlight target deformation. Transfected macrophage attempts to bite DAAMP in half with second untransfected macrophage on the other end, whose presence is implicated by local deformations on the side of the particle not in contact with the transfected cell. Time stamp: min:s. Scale bar, 5 μm.

**Supplementary Movie 7. Contractile activity involved in phagocytic conflict marked by myosin-II.** Maximum intensity projections of RAW macrophages transfected with EGFP-NMMIIA (gray) challenged with DAAM-particles (9 μm,1.4 kPa) (blue). Time stamp: min:s. Scale bar, 5 μm.

**Supplementary Movie 8. Contractile activity leads to sudden forfeit of phagocytic target.** LLSM maximum intensity projections of RAW macrophage expressing mEmerald-Lifeact (gray) challenged with DAAM-particles (9 μm,1.4 kPa) (blue). Transfected cell attempts to internalize second DAAMP target leading to meal sharing event with second, untransfected cell. Dramatic biting of the DAAMP in two leads to the sudden forfeit of the phagocytic target. Merged images (left) with single DAAMP channel in gray (right) to highlight target deformations. Time stamp: min:s. Scale bar, 5 μm.

**Supplementary Movie 9. Contractile activity leads to sudden completion of phagocytic internalization.** LLSM maximum intensity projections of RAW macrophage expressing mEmerald-Lifeact (gray) ingesting DAAM-particles (9 μm,1.4 kPa) (blue). Contractile activity on the DAAMP leads to sudden “popping” of target toward the cell to complete ingestion. Concentrated F-actin ring appears to lag behind this event. Merged images (left) with single DAAMP channel in gray (right) to highlight target displacement. Time stamp: min:s. Scale bar, 5 μm.

**Supplementary references**


1. Vorselen, D. *et al.* Microparticle traction force microscopy reveals subcellular force exertion patterns in immune cell–target interactions. *Nat. Commun.* **11**, 20 (2020).

2. Lankton, S. & Tannenbaum, A. Localizing region-based active contours. *IEEE Trans. image Process.* **17**, 2029–2039 (2008).
